# CD71^+^ erythroid cells exacerbate HIV-1 infection by reactive oxygen species and trans-infect HIV to CD4^+^ T cells

**DOI:** 10.1101/622712

**Authors:** Afshin Namdar, Garett Dunsmore, Petya Koleva, Shima Shahbaz, Juan Jovel, Stan Houston, Shokrollah Elahi

## Abstract

CD71^+^ erythroid cells (CECs) have a wide range of immunomodulatory properties but their potential role in HIV has never been investigated before. Here, we demonstrate that CECs are abundant in the human cord blood, placental tissue and peripheral blood of pregnant mothers. We found that CECs exacerbate HIV-1 infection/replication when co-cultured with CD4^+^ T cells; and that pre-exposure of CD4^+^ T cells to CECs make them more permissible to HIV-infection. Our observations indicate how interactions of CECs with CD4^+^ T cells via reactive oxygen species (ROS)-dependent mechanism results in the upregulation of NF-kB, which affects the cell cycle machinery to facilitate HIV-1 replication. We found the complement receptor-1 (CD35) and the Duffy antigen receptor for chemokines (DARC) as potential HIV-target molecules are expressed significantly higher on CECs compared to mature red blood cells. However, blocking CD35 or DARC did not inhibit HIV-1 trans-infection to uninfected CD4^+^ T cells. We demonstrate that CECs bind to HIV-1 via CD235a and subsequently trans-infect the virus to uninfected CD4^+^ T cells. In addition, we found significant abundance of CECs in the blood of HIV-1 infected and anemic subjects, which enhanced HIV infection/replication in autologous CD4^+^ T cells similar to what we observed for the cord blood and placenta-derived CECs. In agreement, a positive correlation between the frequency of CECs with the plasma viral load in HIV-1 infected antiretroviral therapy naïve individuals was observed. In addition, we found that CECs even in the presence of Tenofovir, can trans-infect HIV-1 to CD4^+^ T cells. Our studies provide a novel insight into the role of CECs in HIV pathogenesis as potential contributing cells for viral persistence in the presence of antiretroviral therapy.

**Author summary:** Despite current antiretroviral therapy, HIV-1 persists in a small pool of infected cells. A better understanding of HIV-reservoirs and influence of other non-immune cells on HIV-1 replication and transmission is a pre-requisite to the development of HIV-eradication strategies. Immature red blood cells (CD71^+^ erythroid cells) are physiologically abundant in newborns, cord blood, placenta and blood of pregnant women, with a wide range of immunological properties. This study demonstrates that these cells not only enhance HIV-1 infection/replication by reactive oxygen species in HIV-target cells (CD4^+^ T cells) but also bind to HIV and trans-infect the virus to the target cells in the presence of Tenofovir, an HIV drug.

We found that these immature red blood cells are abundant in the blood of HIV-patients and anemic individuals. In addition, we observed a positive correlation between the levels of plasma viral load with the frequency of immature red blood cells in HIV-infected individuals. Therefore, our studies discover a novel role for these immature red blood in HIV pathogenesis, which encourages efforts to target these cells as adjuncts of current treatment strategies.

## Introduction

Red blood cells (RBCs) are the most abundant cells in humans (20-30 trillion) and normally generated in the bone marrow (1). Traditionally, the main function of these cells has been considered as gas transporters (O_2_, CO_2_) and preservation of systemic acid/base equilibrium (2). However, in recent years mounting evidence indicate a direct and important role for these cells in innate immunity and inflammation(3, 4). In addition to mature RBCs, a wide range of immunomodulatory functions have been assigned to their immature counterparts (5–8). One important immunomodulatory feature of human RBCs is their tendency to bind to a wide range of chemokines. The Duffy antigen receptor for chemokines (DARC) is an important binding locus, which acts as a sink for IL-8 and interacts with other chemokines(3, 4). Antioxidant property of RBCs is another major immunomodulatory role for these cells, which is essential for their function and integrity. In general, released reactive oxygen species (ROS) by neutrophils and macrophages are taken up by RBCs and get neutralized by the cytosolic antioxidant system (9). However, recent studies indicate that RBCs themselves also generate endogenous ROS(9) and interestingly their immature counterparts mediate immunomodulatory functions via the generation of ROS (10, 11). Immature erythroid cells are typically seen in the periphery of fetuses and newborns while they are absent or in low frequency in the blood of healthy adults(3, 5). It is worth noting that under certain physiological or pathological conditions, immature red blood cells can be seen in the periphery of adults(11–13). For instance, conditions such as anemia, pregnancy, chronic infection and late stage cancer result in a process called extramedullary erythropoiesis in liver and/or spleen, which results with the presence of immature red blood cells in the periphery(6, 10, 11, 14).

Recently, we discovered that immature RBCs (**C**D71^+^ **E**rythroid **C**ells (CECs)) are present in impressively high numbers in neonatal mice spleens and human cord blood. They co-express the transferrin receptor (CD71) and the erythroid lineage marker (TER119) in mice but CD71 and CD235a in humans (15). Additionally, we found that CECs are abundant in the placenta tissue and expand in the peripheral blood of pregnant women (10, 12). We and others have shown CECs have a wide range of immunomodulatory functions and suppress both the innate and adaptive immune responses (6–8, 11, 15). In light of previous reports that HIV-1 through DARC and complement receptor-1 CR1 (CD35) binds to RBCs(16), we speculated that CECs may also express CD35 and DARC. Subsequently, CECs by binding to HIV can impact viral replication and transmission.

HIV pathogenesis in early life, when CECs are physiologically abundant, is defined by rapid CD4^+^ T cell decline, higher plasma RNA levels, accelerated progression to AIDS and death compared to adults(17, 18). Often these infections occur via mother-to-child transmission (MTCT) *in utero*, intrapartum, or post-partum (breastfeeding)(19).

Anemia is another important factor that may influence HIV pathogenesis. It is associated with the abundance of CECs in the peripheral blood. Although the role of anemia in HIV pathogenesis is complex and multifactorial(20, 21), it is a common feature of HIV-related disease and has been uniformly demonstrated to be an independent predictor of morbidity and mortality(21, 22). In addition, anemia can be caused by other pathogens such as malaria, hookworms or micronutrient deficiencies (21, 22). Therefore, due to the physiological expansion of CECs in the periphery of newborns, pregnant mothers, anemic individuals and other pathological conditions (e.g. cancer) it is critical to elucidate their potential effects on HIV transmission and infection/replication.

Here for the very first time, we show that CECs regardless of source (cord blood, placenta, peripheral blood of anemic and HIV patients) mediate the exacerbation of HIV-1 replication in CD4^+^ T cells. Our observations coupled with RNAseq data demonstrate how interactions of CECs with CD4^+^ T cells via ROS affect the cell cycle machinery to facilitate HIV-1 replication. In addition, we demonstrate that CECs compared to mature RBCs express significantly higher levels of both CD35 and DARC, and can trans-infect HIV-1 to uninfected CD4^+^ T cells *in vitro*. Finally, our study indicates that HIV-1 interacts with CD235a and thus CECs can trans-infect uninfected CD4^+^ T cells in the presence of anti-retroviral drug, Tenofovir.

## Results

### The cord blood and placental CECs exacerbate HIV infection in already infected autologous CD4^+^ T cells

In agreement with our previous reports (6, 15), here we show that CECs are physiologically abundant in the human umbilical cord blood, and placental tissues while they are almost absent in the peripheral blood of healthy adults (Fig. 1A and B). Since we have reported that CECs have immunomodulatory properties (6, 8, 15), their impact on HIV-1 infection in CD4^+^ T cells was evaluated in an HIV-1 *ex vivo* infection assay. Cord blood CD4^+^ T cells were isolated and made more permissible to HIV-1 infection by *in vitro* culture with exogenous IL-2 and PHA stimulation (23). Subsequently, CD4^+^ T cells were infected with either the lab-adapted X4-tropic isolate (HIV-1_LAI_) or R5-tropic HIV-1 isolate (HIV-1_JR-CSF_). Isolated autologous CECs at different ratios were added to the infected CD4^+^ T cells following extensive wash to remove extracellular viruses. Viral replication was analyzed by intracellular p24 staining using flow cytometry 3-4 days later. Using these culture conditions, we consistently observed that CECs significantly enhanced HIV infection in CD4^+^ T cells with both X4-tropic (Fig. 1C and D) and R5-tropic HIV-1 viruses (Fig. 1E and F). CECs mediated enhanced HIV-1 infection in CD4^+^ T cells was dose dependent for both X4-tropic and R5-tropic viral isolates, respectively (Fig. 1D and F). We found that CECs not only significantly increased the number of infected CD4^+^ T cells (Fig. 1C-F) but also the number of virus per cell was significantly greater as shown by the intensity of p24 expression (Fig. 1G and SI Appendix, Fig. S1A and S1B). Consistent with activated CD4^+^ T cells, we found that CECs increased HIV-1 infection in non-activated CD4^+^ T cells (Fig. 1H and I). We also found that placenta-derived CECs similar to the cord blood, significantly increased HIV-1 infection in autologous CD4^+^ T cells (Fig. 1J and K). Since non-activated cord blood CD4^+^ T cells do not express substantial levels of CCR5 compared to CXCR4 (SI Appendix, Fig. S1C) and CECs enhanced replication of both viral isolates, we decided using X4-tropic viral isolate for the subsequent studies.

**Fig. 1.**
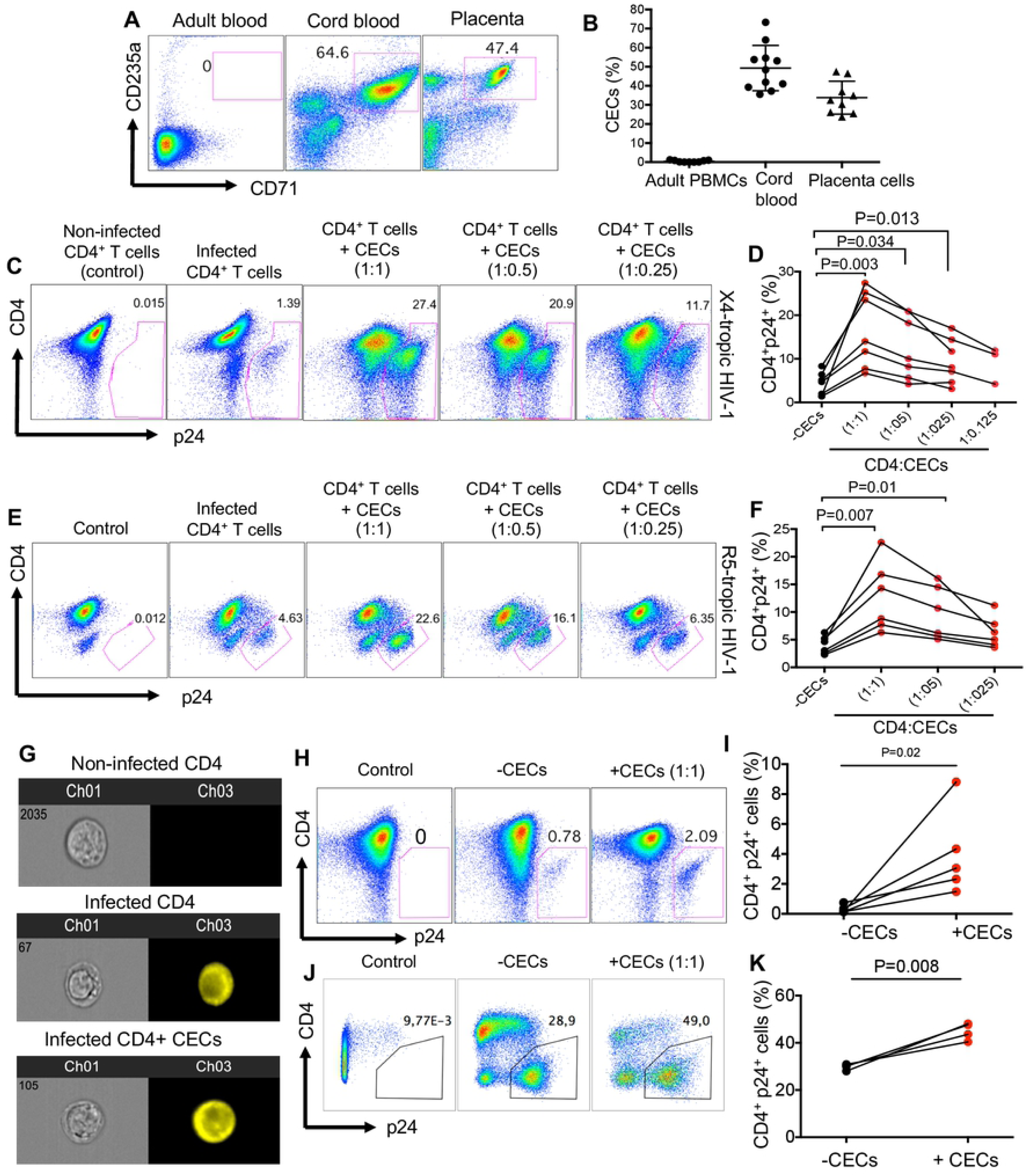
The cord blood and placenta CECs exacerbate HIV infection in already infected autologous CD4^+^ T cells. (**A**). Representative plots showing frequency of CECs in the adult peripheral blood mononuclear cells (PBMCs) versus the cord blood mononuclear cells (CBMCs) and placenta. (**B**) Cumulative data comparing the percentages of CECs in adult PBMCs versus CBMCs and placental tissues. (**C**) Representative flow cytometry plots showing activated CD4^+^ T cells from the cord blood following infection with X4-tropic (HIV-1 _LAI_) isolate in the absence or presence of CECs. (**D**) Cumulative data showing percentages of activated CD4^+^ T cells after infection with X4-tropic isolate and co-cultured with CECs at different ratios. (**E**) Representative flow cytometry plots showing activated CD4^+^ T cells from cord blood following infection with R5-tropic (HIV-1 _JR-CSF_) isolate in the absence or presence of CECs. (**F**) Cumulative data showing percentages of activated CD4^+^ T cells after infection with R5-tropic isolate in the presence or absence of CECs at different ratios. (**G**) Representative images showing intensity of p24 in CD4^+^ T cells in the absence or presence of CECs. (**H**) Representative flow cytometry plots showing inactivated CD4^+^ T cells from the cord blood following infection with X4-tropic isolate in the absence or presence of CECs. (**I**) Cumulative data showing percentages of non-activated CD4^+^ T cells after infection with X4-tropic isolate in the absence or presence of CECs. (**J**) Representative flow cytometry plots showing activated CD4^+^ T cells from placenta following infection with X4-tropic isolate in the absence or presence of placenta CECs. (**K**) Cumulative data showing percentages of activated and infected CD4^+^ T cells from placenta with X4-tropic viral isolate in the absence or presence of CECs. The number of infected cells was quantified by intracellular viral p24 antigen staining using flow cytometry on day 4 post infection. Each point represents a cord blood or a placenta.

### Pre-exposure of CD4^+^ T cells to autologous cord blood CECs enhance their infectivity to HIV-1 infection

Activated CD4^+^ T cells were co-cultured with CECs (1:1 ratio) overnight, then CD4^+^ T cells were isolated and infected with HIV-1_LAI._ We observed that pre-exposure of CD4^+^ T cells to CECs significantly enhanced their infectivity to HIV-1 (Fig. 2A and B). To confirm if this was the case for non-activated CD4^+^ T cells, similar overnight co-cultures were performed. Likewise, we observed pre-exposure of resting CD4^+^ T cells to CECs significantly enhanced their susceptibility to HIV-1 infection (Fig. 2C and D).

**Fig. 2.**
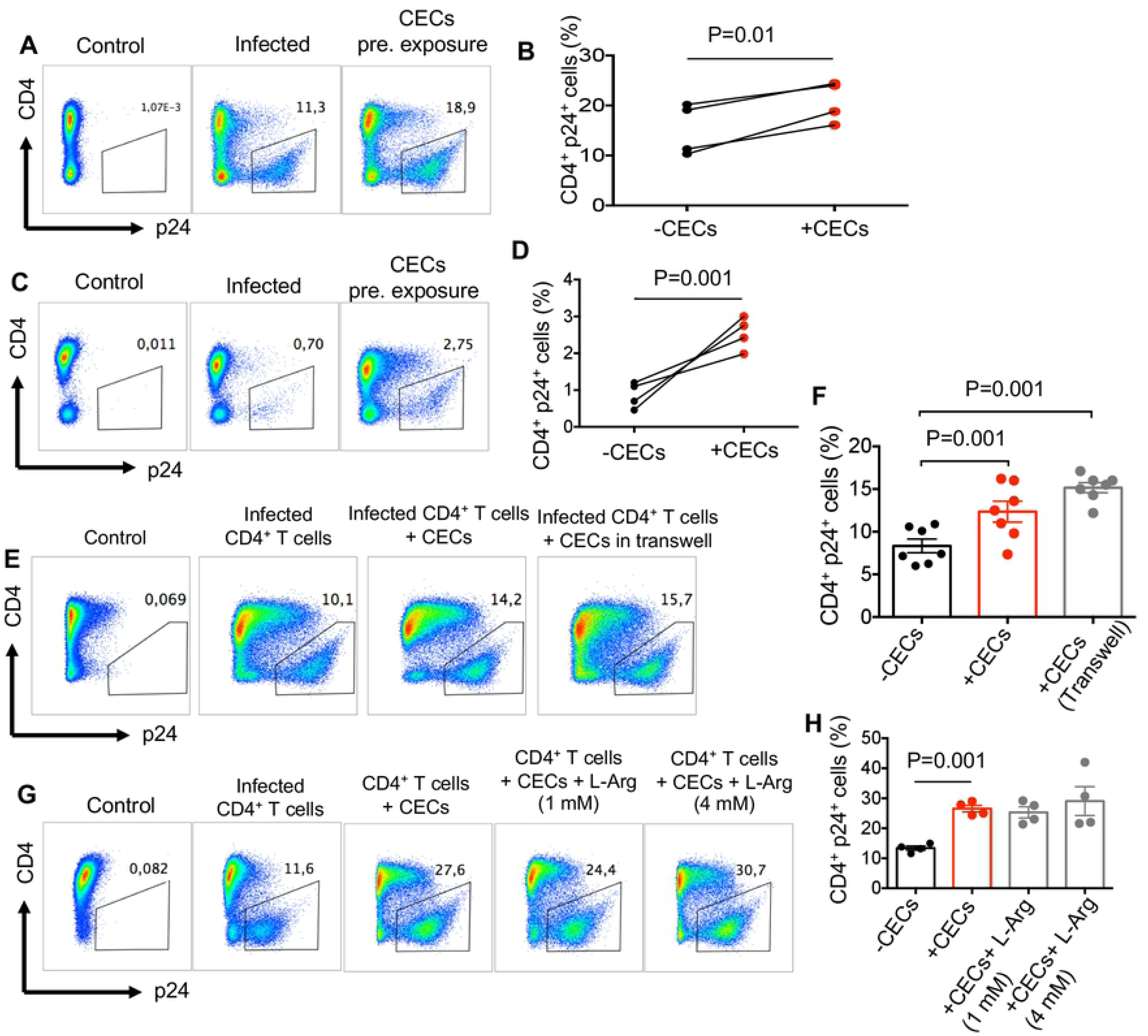
Pre-exposure of CD4^+^ T cells to autologous cord blood CECs make them more permissible to HIV-1 infection. (**A**) Representative flow cytometry plots showing pre-exposed activated CD4^+^ T cells to CECs prior to infection with X4-tropic isolate versus non-exposed activated CD4^+^ T cells to CECs (**B**) Cumulative data showing percentages of p24 in activated CD4^+^ T cells when either pre-exposed to CECs prior to infection with X4-tropic isolate or were not exposed to CECs. (**C**) Representative flow cytometry plots showing pre-exposed non-activated CD4^+^ T cells to CECs prior to infection with X4-tropic isolate versus non-exposed non-activated CD4^+^ T cells to CECs (**D**) Cumulative data showing percentages of p24 in non-activated CD4^+^ T cells when either pre-exposed to CECs prior to infection with X4-tropic isolate or were not exposed to CECs. (**E**) Representative flow cytometry plots showing infection with X4-tropic isolate in activated CD4^+^ T cells when co-cultured with CECs versus trans-well system (**F**) Cumulative data showing percentages of infected CD4^+^ T cells with X4-tropic when co-cultured with CECs versus trans-well system. (**G**) Representative flow plots showing infection with X4-trpoic isolate in activated CD4^+^ T cells in the presence of CECs and/or L-arginine (L-Arg) at 1 and 4mM. (**H**) Cumulative data showing percentages of infected CD4^+^ T cells with HIV in the presence or absence of CECs and/or L-arginine.

### CECs via soluble mediators enhance HIV-1 infection/replication in CD4^+^ T cells

We further decided to determine whether enhanced HIV-1 infection in CD4^+^ T cells required cell-cell interactions. By using a Transwell system we observed that increased HIV-1 infection in CD4^+^ T cells does not necessarily depend on cell-cell interactions (Fig. 2E and F). Previously, we had demonstrated that CECs mediate immunosuppression via enzymatic activity of arginase-2 (6, 8, 12, 15). In addition, high level of arginase activity and subsequent _L_-arginine deprivation has been associated with dysfunctional T cells and increased viral load in HIV-patients (24). Thus, we hypothesized that arginaze-2 activity by CECs may enhance HIV-1 infection/replication in CD4^+^ T cells. In contrast, we found that overriding the enzymatic activity of arginase-2 by _L_-arginine supplementation did not abrogate the enhanced HIV-1 infection in CD4^+^ T cells when co-cultured with CECs (Fig. 2G and H). Despite the fact that CECs constitutively secrete TGF-β (25), we found this cytokine had no effects on CECs-mediated enhanced HIV-infection/replication in CD4^+^ T cells (data not shown). These data suggest that CECs possibly by secreting other soluble factors promote HIV-1 infection/replication.

### CECs via oxygen containing compounds modulate the expression of genes associated with increased CD4^+^ T cells infectivity

We implemented RNA sequencing (RNA-seq) to analyze the transcriptome of HIV-1 infected CD4^+^ T cells alone or when co-cultured with CECs. When hierarchical clustering was conducted on Euclidian distances between samples, CD4^+^ T cells exposed to CECs clearly showed a different gene expression profile compared to CD4^+^ T cells that were not co-cultured with CECs (SI Appendix Fig. S1D). Those results were recapitulated in principal component analysis (PCA) on the Euclidian distances between samples (SI Appendix Fig. S1E). In essence, we found the transcriptome profile of infected CD4^+^ T co-cultured with CECs was clearly separated from the infected CD4^+^ T cells in the absence of CECs (Fig. 3A). Gene ontology of biological process for the transcriptome profile revealed upregulation of cellular response to oxygen-containing compounds and upregulation of NF-kB signaling in CD4^+^ cells co-cultured with CECs (Fig. 3B). In contrast, negative regulation of cellular protein metabolic/catabolic process was evident in highly downregulated genes (Fig. 3C). In total, 1011 genes were highly upregulated (≥ 2-fold, FDR ≤ 0.05) and 1910 genes downregulated (≥ 2-fold, FDR ≤ 0.05) in co-cultured infected CD4^+^ T cells compared to CD4^+^ T cells alone. We selected approximately 80 highly upregulated and downregulated genes (SI Appendix, Fig. S1F). The biological functions of most of these genes in regards to HIV-1 replication/T cells proliferation are unknown. However, among these, we identified 16 highly upregulated genes in CD4^+^ T cells when co-cultured with CECs (Fig. 3D), which can be associated with CECs mediated enhanced HIV-1 infection. The tissue transglutaminase (tTg) gene was the most upregulated gene (> 12 folds) followed by AQP9 (aquaporin 9). Aquaporins (AQPs > 12 folds) are channel proteins widely present in living cells to facilitate the transport of water and certain neutral solutes across biological membranes. The third highly upregulated gene was MYOF (> 10 folds), which is a membrane-associated protein involved in both caveolin and clathrin-mediated endocytosis pathways along with membrane fusion after damage (26). It has been reported that upregulation of AQP9 is required as ROS scavenger to prevent T cells apoptosis (27). Other highly upregulated genes were ASAP1 (ArfGAP with SH3 Domain, Ankyrin Repeat and PH Domain 1), KYNU (Kynurenine), LRRK2 (The leucine-rich repeat kinase 2), TCL1A (The T cell leukemia-lymphoma 1 TCL1), TJP2 (tight junction protein 2), BCLAF1 (Bcl-2-associated transcription factor 1), RHBDD1 (The rhomboid domain containing 1), MEF2C (the myocyte enhancer factor-2C), IDO1 (indoleamine 2,3-dioxygenase 1), POGZ (pogo transposable element-derived protein with zinc finger domain) and inhibitor of nuclear factor kappa B kinase subunit B IKBKB (Fig. 3D). Upregulation of NF-kB gene expression was confirmed by qPCR, as shown in Fig. 3E, NF-kB mRNA was 3 folds higher in HIV-1 infected CD4^+^ T cells when co-cultured with CECs relative to HIV-1 infected CD4^+^ T cells alone.

**Fig. 3.**
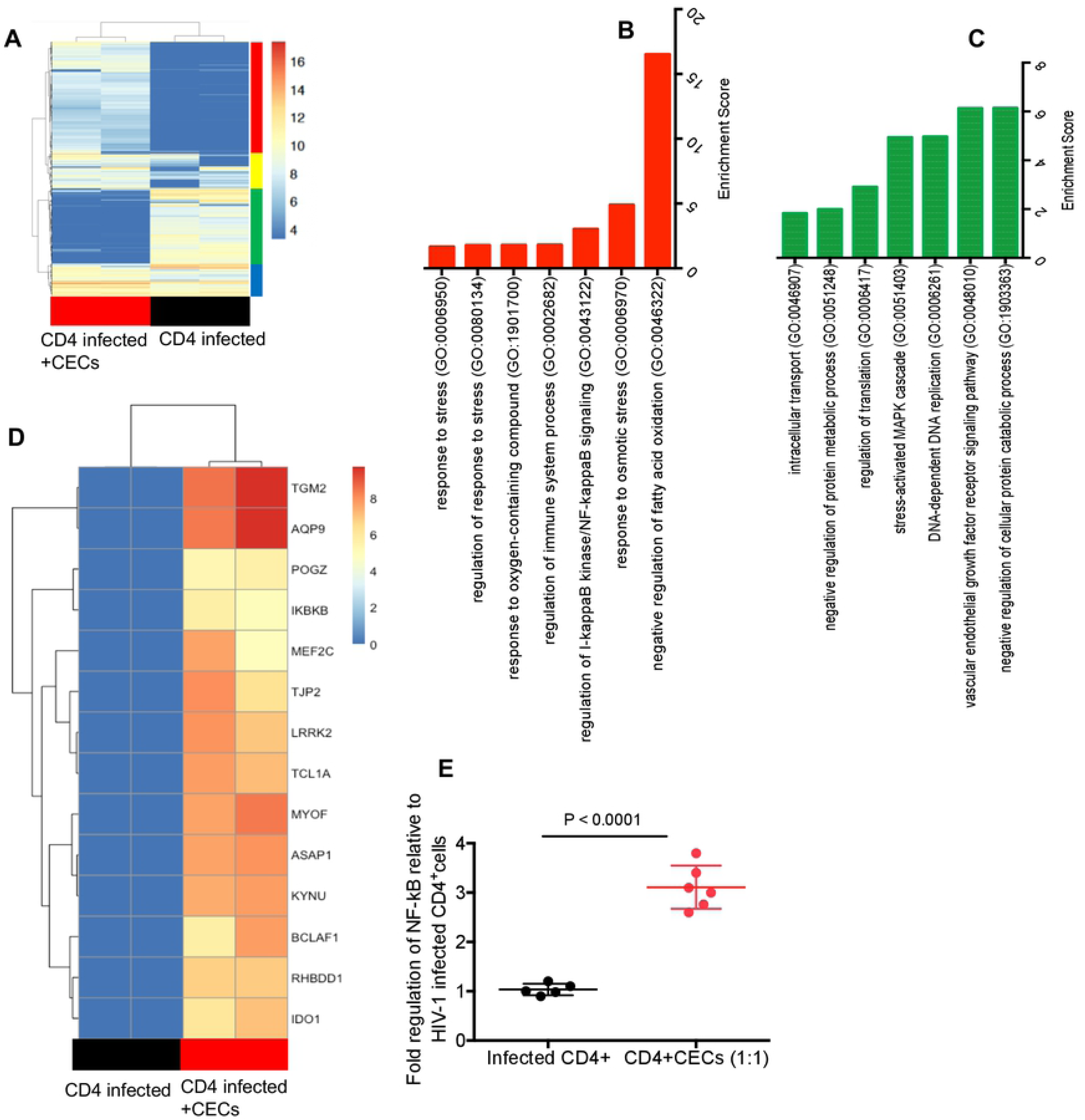
CECs via oxygen containing compounds modulate the expression of genes associated with increased CD4^+^ T cells infectivity. (**A**) Transcriptome of infected CD4^+^ T cells in the absence and presence of CECs is shown using heatmap. (**B**) Gene ontology of biological process for the transcriptome profile of upregulated genes in CD4^+^ T cells co-cultured with CECs. (**C**) Gene ontology of biological process for the transcriptome profile of down-regulated genes in CD4^+^ T cells in the presence of CECs. (**D**) Most upregulated genes associated with enhanced HIV-infection in CD4^+^ T cells. (**E**) NF-kB gene expression was confirmed in CD4^+^ T cells when co-cultured with CECs. The underlying source data are provided as a Source Data file.

### CECs enhance HIV infection via ROS

It has been reported that RBCs secrete endogenous ROS (28) and the hydrogen peroxide produced by these cells exhibits various immunological functions (29). More importantly, gene ontology of biological process of the transcriptome profile of co-cultured CD4^+^ T cells with CECs, revealed upregulation of cellular response to oxygen-containing compounds and NF-kB signaling (SI Appendix, Fig. S2A). Therefore, we performed qPCR on CECs for the expression of NOX family genes which comprises of seven paralogous including NOX1-5 and dual oxidase (DUOX1/2) (30). Interestingly, we observed a prominent gene expression of NOX2 in CECs (Fig. 4A), while NOX paralogous (NOX 1, 3, 4, 5, DUOX1, and 2) were undetectable. We further compared ROS production by RBCs (CD235a) versus CECs either from the cord blood or placenta. We found that CECs compared to their mature counterparts produce significantly higher levels of ROS (Fig. 4B and C) and CECs from the placenta express significantly higher levels of ROS compared to their counterparts in the cord blood (Fig. 4D and E). In agreement with our previous report, we observed lower ROS expression by CECs from the cord blood and placenta of inflammatory bowel disease (IBD) patients compared to healthy controls (10) (Fig. 4F and G). This indicates a dichotomy in ROS production by CECs from IBD patients compared to the healthy. However, there was no significant difference in mRNA expression levels for arginase-2 in CECs isolated from the cord blood or placenta of IBD versus healthy samples (SI Appendix, Fig. S2B and 2C). We further aimed to determine the role of NADPH oxidases (NOX) complex, which is involved in ROS generation. Due to the dichotomy in ROS production by CECs from IBD patients compared to healthy donors, we decided to perform HIV-1 infection assays using autologous CD4^+^ T cells and CECs from the cord blood of IBD patients.

**Fig. 4.**
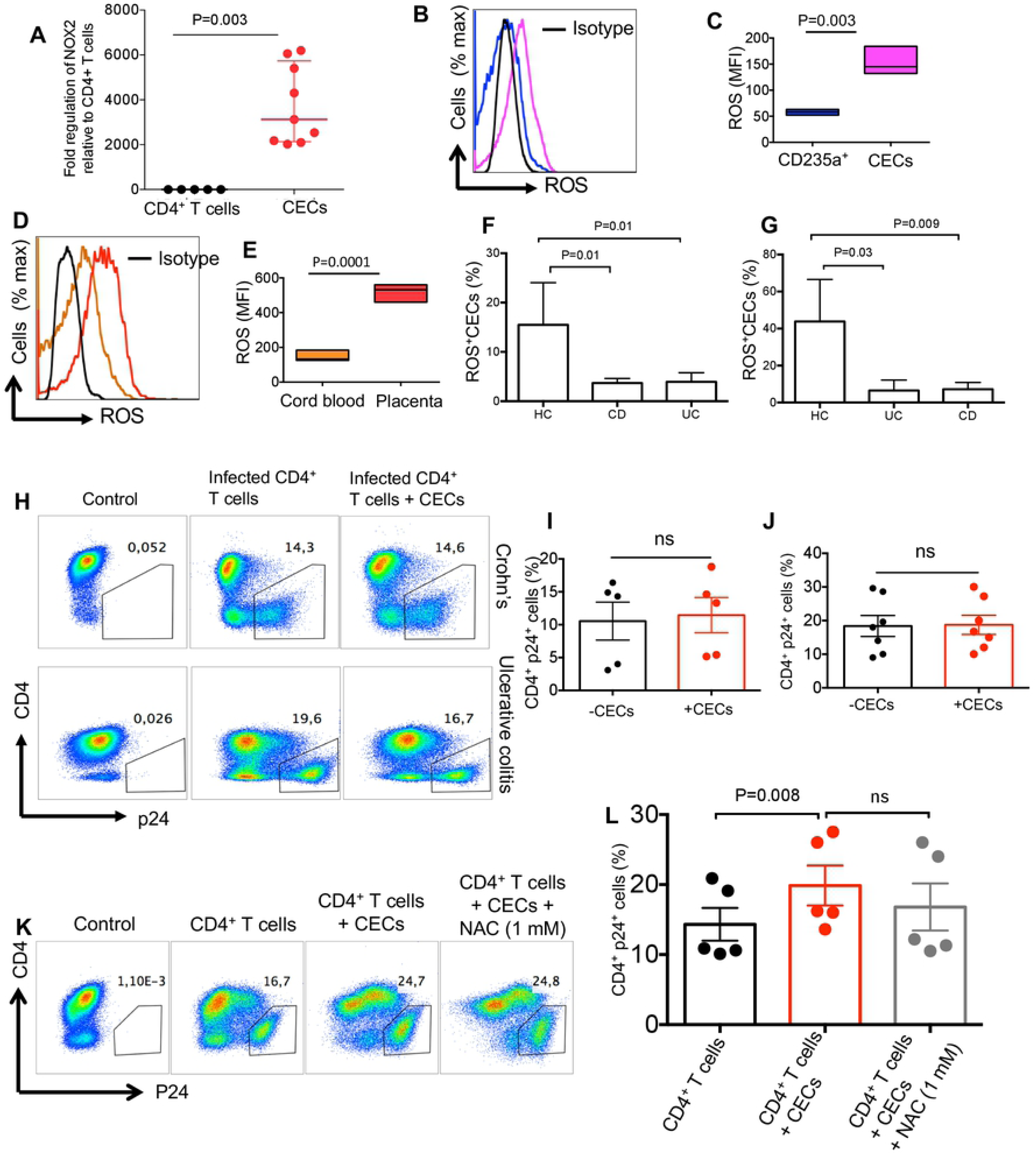
CECs enhance HIV infection through ROS. (**A**) NOX2 gene expression in CECs versus CD4^+^ T cells. (**B**) Representative histogram showing ROS expression (RBCs blue line, CECs purple line) or isotype control (black line). (**C**) The MFI of ROS among RBCs (CD235a^+^) and CECs. (**D**) Representative histogram showing ROS expression in CECs from the cord blood (orange line) or the placenta (red line) and isotype control (black line). (**E**) The Mean Fluorescence intensity (MFI) of ROS among the cord blood or placenta CECs. (**F**) Cumulative data showing % ROS^+^ CECs in the cord blood or (**G**) the placenta of healthy controls (HC) versus patients with Crohn’s disease (CD) or Ulcerative colitis (UC). (**H**) Representative flow cytometry plots showing HIV-infection in activated CD4^+^ T cells in the presence or absence of cord blood CECs. (**I**) Cumulative data showing %p24 in activated CD4^+^ T cells in the presence or absence of CECs from the cord blood of newborns to CD or (**J**) UC mothers. (**K**) Representative flow cytometry plots showing HIV-infection in the cord blood CD4^+^ T cells in the presence or absence of CECs and N-acetyl cysteine (NAC) (**L**) Cumulative data showing %p24 in activated CD4^+^ T cells in the presence/absence of CECs and/or NAC.

Interestingly, we found lack of enhanced HIV replication in autologous CD4^+^ T cells by CECs obtained from the cord blood of IBD donors (Fig. 4H-J). Next, we assessed whether down-regulation of ROS activity by ROS inhibitors could mitigate HIV infection in CD4^+^ T cells. To evaluate this, HIV infected CD4^+^ T cells in the presence of CECs were exposed to two types of ROS inhibitors; N-acetyl cysteine (NAC) and Apocynin (Apo). The universal ROS blocker (NAC), failed to inhibit the enhanced viral replication by CECs in CD4^+^ T cells even at 1mM concentration (Fig. 4K and L). In contrast, Apo, which is the NADPH-dependent ROS inhibitor, abrogated the enhanced HIV-1 infection/replication in CD4^+^ T cells by CECs (Fig. 5A and B). This observation suggested that CECs might release mitochondrial ROS, which was confirmed by MitoSOX indicator (Fig. 5C-F). In addition, we confirmed that CECs from either the cord blood (Fig. 5C) or the placenta (Fig. 5D) had higher mitochondrial superoxide compared to RBCs (Fig. 5E).

**Fig. 5.**
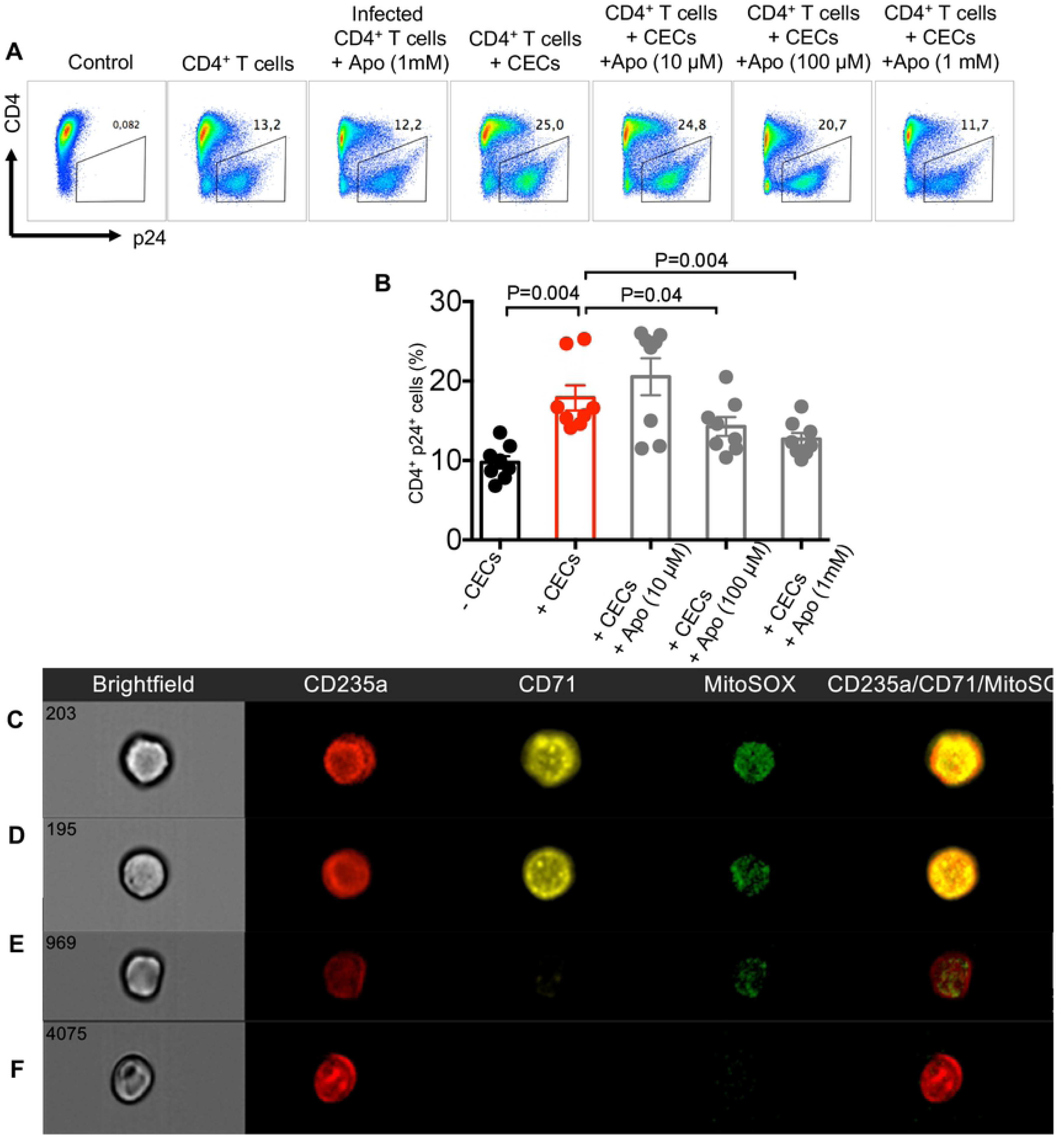
Apocynin abrogates CECs mediated HIV-infection by ROS. (**A**) Representative flow cytometry plots showing HIV-infection in the cord blood CD4^+^ T cells in the presence or absence of CECs and Apocynin (Apo) (**B**) Cumulative data showing %p24 in activated CD4^+^ T cells in the presence/absence of CECs and/or Apo. (**C**) Representative of images collected using an Amnis Imagestream Mark II showing MitoSOX in the cord blood or (**D**) placenta CECs and (**e**) RBCs compared to (**f**) ROS isotype control.

### CECs express significantly greater levels of DARC and CD35 compared to RBCs

The role of DARC on RBCs in mediating trans-infection of HIV-1 to the target cells has already been documented (16). Therefore, we decided to determine whether CECs similar to their older siblings express DARC. Interestingly, we observed nearly 100% of CECs express DARC compared to approximately 20% of RBCs (Fig. 6A-C). Additionally, the intensity of DARC was significantly higher on CECs compared to RBCs (Fig. 6D).

**Fig. 6.**
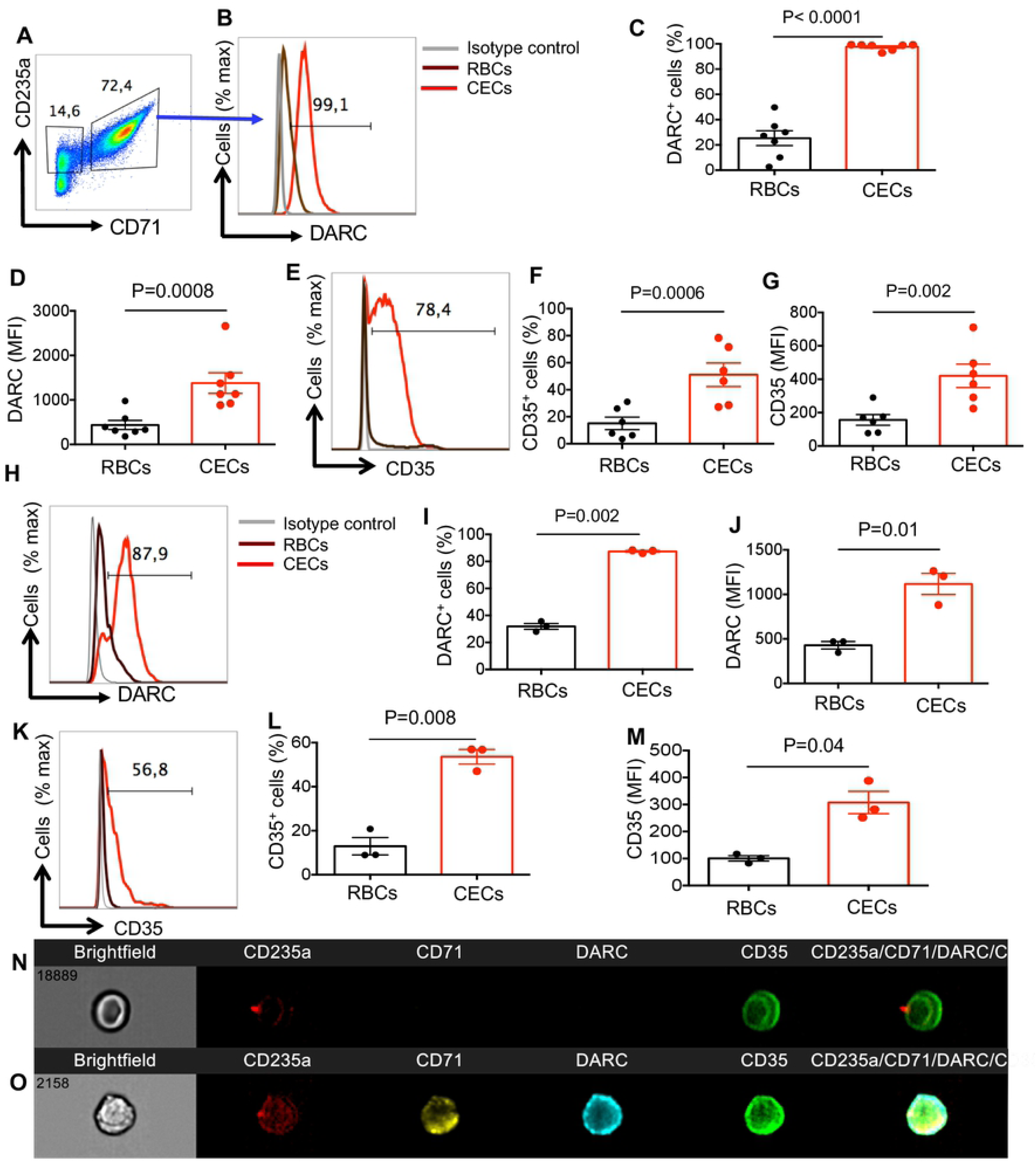
DARC and CD35 are highly expressed on CECs. (**A**) Representative flow cytometry plots showing gating strategy for CECs (CD71^+^CD235a^+^) versus RBCs (CD235a^+^). (**B**) Representative histogram showing DARC expression on CECs from the cord blood (red line) or RBCs (brown line) and isotype control (black line). (**C**) Cumulative data showing % DARC expressing or (**D**) the MFI of DARC expression on RBCs versus CECs from the cord blood. (**E**) Representative histogram showing CD35 expression on CECs from the cord blood (red line) or RBCs (brown line) and isotype control (black line). (**F**) Cumulative data showing % CD35 expressing or (**G**) the MFI of CD35 expression on RBCs versus CECs from the cord blood. (**H**) Representative histogram showing DARC expression on CECs from the placenta (red line) or RBCs (brown line) and isotype control (black line). (**I**) Cumulative data showing % DARC expressing or (**J**) the MFI of DARC expression on RBCs versus CECs from the placenta. **(K**) Representative histogram showing CD35 expression on CECs from the placenta (red line) or RBCs (brown line) and isotype control (black line). (**L**) Cumulative data showing % CD35 expressing or (**M**) the MFI of CD35 expression on RBCs versus CECs from the placenta. (**N**) Representative of images collected using an Amnis Imagestream Mark II showing DARC and CD35 expression on RBCs or (**O**) CECs from the cord blood.

Another molecule of interest was CD35 (CR-1), which its opsonised role for HIV particles via HIV/anti-HIV immune complexes and complement factor C3b binding has already been demonstrated (31). Therefore, we measured the expression level of CR-1 on both RBCs and CECs. Interestingly, we found substantial surface expression of CR-1 on CECs, which was significantly higher in terms of percentage and intensity compared to RBCs (Fig. 6E-G). Similar pattern of expression was observed for both DARC (Fig. 6H-J) and CD35 (6K-M) on placental CECs. Finally, image stream analysis confirmed the higher expression of DARC and CD35 on CECs versus RBCs (Fig. 6N and O). Thus, our observations indicate prominent expression of DARC and CD35 on CECs.

### CECs mediate trans-infection of HIV-1 to CD4^+^ T cells

To determine whether CECs mediate HIV-1 trans-infection to CD4^+^ T cells, we incubated CECs with HIV-1 (exactly similar to the methods used for infecting CD4^+^ T cells). Then extensively washed and co-cultured with autologous uninfected CD4^+^ T cells. We found that pre-exposure of CECs to HIV-1 resulted in carriage of the virus and trans-infection of CD4^+^ T cells when analyzed 4 days post-coculture (Fig. 7A and B). Interestingly, the infection rate in trans-infected CD4^+^ T cells was slightly lower to the infection rate of CD4^+^ cells when co-cultured with CECs but significantly higher than infected CD4^+^ T cells in the absence of CECs (Fig. 7B). Furthermore, we decided to evaluate the effects of DARC and CD35 blockade on HIV trans-infection by CECs since such role for RBCs has been reported (16, 31). First of all, we showed that addition of anti-CD35 antibody, CCL-5 (to compete for DARC) and/or their combination had no significant effects on HIV infection/replication in CD4^+^ T cells alone (Fig. 7C and D). Then we investigated the effects of anti-CD35 antibody, recombinant CCL-5 (rCCL-5) and their combination on infected CD4^+^ T cells when co-cultured with CECs. As expected, addition of CECs enhanced HIV infection in CD4^+^ T cells however blocking CD35 or addition of rCCL-5 had no significant effects on the infection rate but their combination slightly reduced HIV-infection (Fig. 7E and F). Next, we decided to interfere with the interaction of HIV with its potential targets (DARC and CD35) on CECs by pre-incubating CECs with anti-CD35 antibody, rCCL-5 or their combination for an hr prior to the viral exposure. Following incubation, HIV-1 was added to CECs in the presence of anti-CD35 antibody and rCCL-5. Interestingly, we found that pre-exposure to rCCL-5, anti-CD35 antibody or their combination had no significant effects on the ability of CECs to trans-infect CD4^+^ T cells using magnetofection (Fig. 7G and H). However, their combination slightly reduced HIV-infection without using magnetofection approach (Fig.7I and J). To exclude possible interference of serum in culture media on the activity of anti-CD35 antibody, we performed CD35 blocked in the absence of serum. However, we did not observe any reduction in trans-infection of HIV-1 to CD4^+^ T cells (SI Appendix, Fig. S2D and 2E). Finally, we found that pre-exposure of CECs to HIV-1 also trans-infect the virus to non-activated CD4^+^ T cells, though in lower rate as predicted (SI Appendix, Fig. S2F). These observations indicate that CECs were able to trans-infect HIV-1 to uninfected CD4^+^ T cells even in the presence of rCCL-5 and/or anti-CD35 antibody or their combination.

**Fig. 7.**
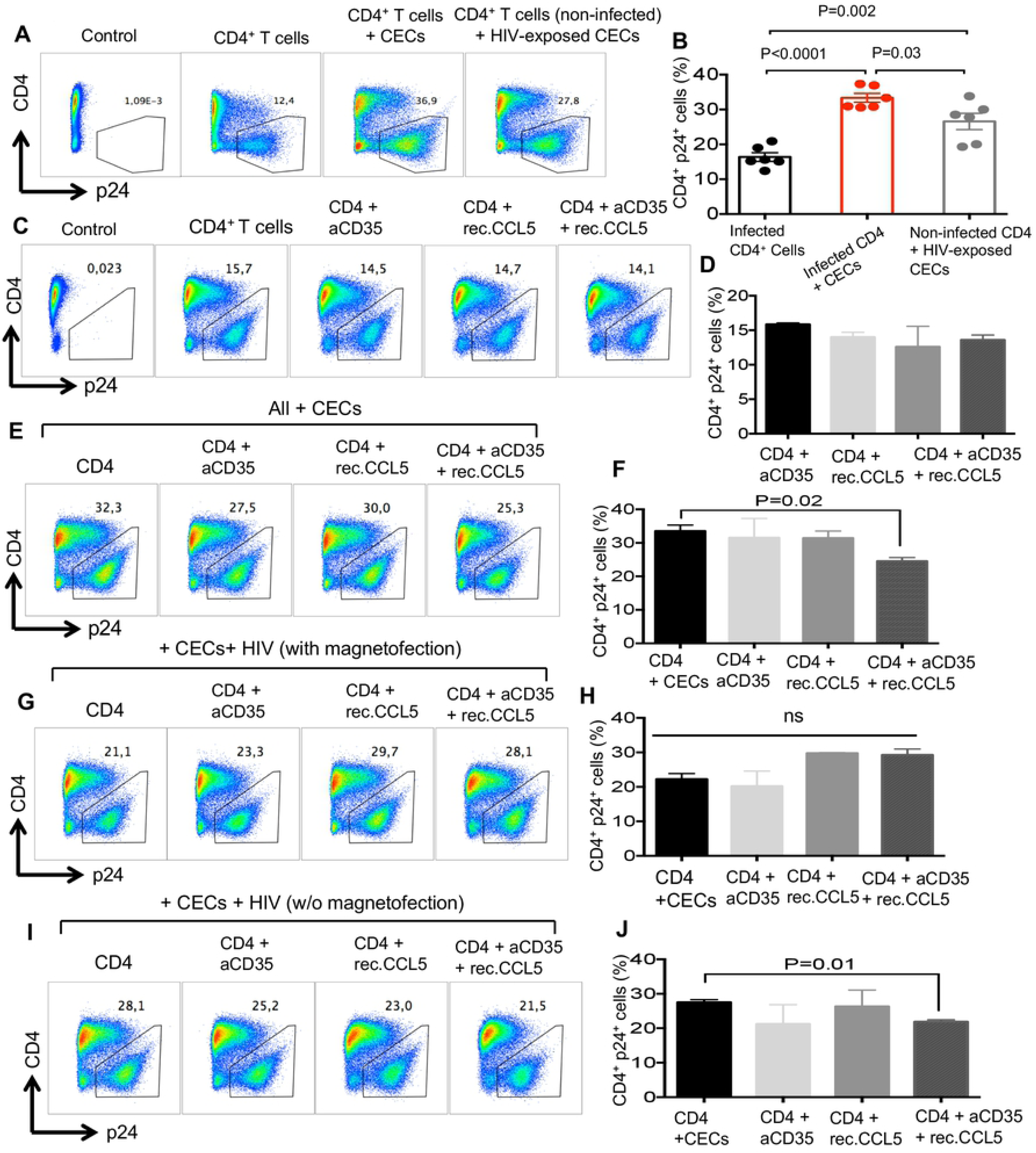
CECs mediate trans-infection of HIV-1 to CD4^+^ T cells. (**A**) Representative flow cytometry plots showing CD4^+^ T cells from the cord blood following infection with X4-tropic isolate in the absence/presence of CECs and also CECs exposed to HIV then co-cultured with non-infected but activated CD4^+^ T cells. (**B**) Cumulative data showing %p24 in CD4^+^ T cells in the absence/presence of CECs and also CECs exposed to HIV then co-cultured with non-infected but activated CD4^+^ T cells. (**C**) Representative flow cytometry plots showing infected CD4^+^ T cells in the presence of anti-CD35 antibody (5µg/ml), CCL-5 (1µM) or their combination (anti-CD35 (5µg/ml) and CCL-5 (1µM)). Cumulative data showing %CD4^+^p24^+^ T cells in the presence of anti-CD35 antibody (5µg/ml), CCL-5 (1µM) or their combination (anti-CD35 (5µg/ml) and CCL-5 (1µM)). (**D**) Representative flow cytometry plots showing infected CD4^+^ T cells in the presence of CECs alone or in the presence of CECs plus anti-CD35 antibody (5µg/ml), CCL-5 (1µM) or their combination (anti-CD35 (5µg/ml) and CCL-5 (1µM)). (**F**) Cumulative data showing %CD4^+^p24^+^ T cells in the presence of CECs alone or in the presence of CECs plus anti-CD35 antibody (5µg/ml), CCL-5 (1µM) or their combination (anti-CD35 (5µg/ml) and CCL-5 (1µM)). (**G**) Representative flow cytometry plots showing infected CD4^+^ T cells in the presence of CECs alone or in the presence of pre-exposed CECs to anti-CD35 antibody (5µg/ml), CCL-5 (1µM) or their combination (anti-CD35 (5µg/ml) and CCL-5 (1µM)) using magnetofection. (**H**) Cumulative data showing %CD4^+^p24^+^ T cells in the presence of CECs alone or in the presence of CECs plus anti-CD35 antibody (5µg/ml), CCL-5 (1µM) or their combination (anti-CD35 (5µg/ml) and CCL-5 (1µM)) using magnetofection. (**I**) Representative flow cytometry plots showing infected CD4^+^ T cells in the presence of CECs alone or in the presence of CECs plus anti-CD35 antibody (5µg/ml), CCL-5 (1µM) or their combination (anti-CD35 (5µg/ml) and CCL-5 (1µM)) without magnetofection (**J**) Cumulative data showing %CD4^+^p24^+^ T cells in the presence of CECs alone or in the presence of CECs plus anti-CD35 antibody (5µg/ml), CCL-5 (1µM) or their combination (anti-CD35 (5µg/ml) and CCL-5 (1µM)) without magnetofection.

### HIV-1 preferentially binds to CD235a on CECs but not RBCs and Tenofovir treated CECs trans-infect HIV to uninfected CD4^+^ T cells

Since blockade of CD35 and addition of rCCL-5 did not abrogate CECs-mediated HIV trans-infection to CD4^+^ T cells, we decided to investigate the interaction of HIV with CECs using a CCR5-tropic GFP marked virus. Using image stream, we observed that HIV appears to be interacting more strongly with CECs (Fig. 8A and B) compared to RBCs (Fig. 8C). Most interestingly, we found co-localization of CD235a with GFP on CECs, which suggests interaction of HIV with CD235a (Fig. 8A and B). However, this was not prominent for RBCs (Fig. 8C). These observations were further supported by significantly higher MFI for CD235a on CECs versus RBCs (Fig. 8D). Moreover, we exposed CECs to the X4-tropic HIV-1 isolate according to our routine infection protocol(23) and 3 days later infection rate in CECs was quantified by measuring p24. Interestingly, a small portion of CECs was positive for p24 (Fig. 8E and F). Although the majority of the cord blood CECs lack nuclei, about 5% of them still have nuclei, which may explain these observations.

**Fig. 8.**
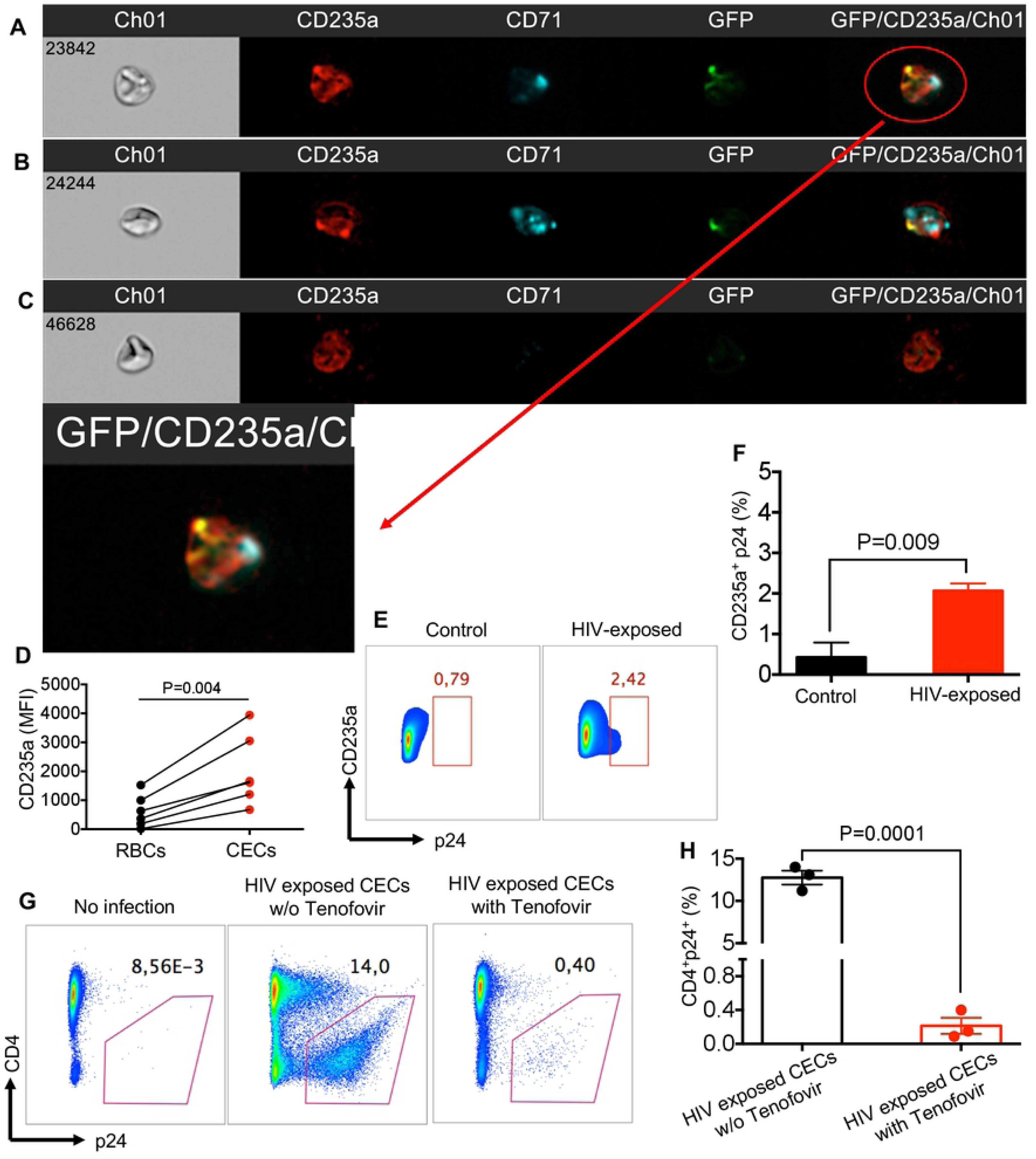
HIV-1 preferentially binds to CECs and CECs-mediated HIV trans-infection occurs in the presence of Tenofovir. (**A and B**) Representative image stream plots showing interaction of CECs with GFP-marked HIV versus (**C**) RBC. (**D**) Cumulative data showing MFI of CD235a on CECs versus RBCs. (**E**) Representative flow cytometry plots showing p24 in HIV-exposed CECs versus control. (**F**) Cumulative data showing %p24 in HIV-exposed CECs versus controls. (**G**) Representative flow cytometry plots showing HIV-trans-infection by CECs to CD4^+^ T cells in the presence and absence of Tenofovir (4.3 ug/ml). (**H**) Cumulative data showing HIV-trans-infection by CECs to CD4^+^ T cells in the presence and absence of Tenofovir.

To further understand the physiological relevance of CECs-mediated HIV-1 trans-infection in HIV-patients on anti-retroviral therapy (ART), CECs were exposed to HIV for 40 min, then washed extensively and incubated with Tenofovir (4.3 µg/ml) for 16 hrs, which is physiologically relevant considering 300 mg/daily recommendation. CECs were then washed and co-cultured with uninfected but activated autologous CD4^+^ T cells for 3 days. Our observations indicated that although Tenofovir significantly reduced HIV-1 infection, it did not eliminate all the bound or possibly internalized viruses in CECs (Fig. 8G and H).

### Increased frequency of CECs in the peripheral blood of HIV-1 infected and anemic individuals

We found CECs become abundant in the blood of HIV-infected individuals versus healthy controls (Fig. 9A and B) and generate ROS (Fig. 9C). Similarly, we observed expansion of CECs in the periphery of anemic individuals (Fig. 9D). Since HIV-induced CECs produce ROS, we reasoned these cells similar to the cord blood CECs might enhance HIV-1 infection in CD4^+^ T cells. In agreement, we found enhanced HIV-1 infection in CD4^+^ T cells when co-cultured with autologous CECs *in vitro* (Fig. 9E). More importantly, we observed a positive correlation between the frequency of CECs with the plasma viral load in HIV-infected and ART-naïve patients (Fig. 9F). We also found that CECs isolated from the blood of anemic individuals enhance HIV-1 infection/replication when co-cultured with autologous CD4^+^ T cells (Fig. 9G and H). Finally, we found that CECs obtained from HIV+ (Fig. 9I) and anemic individuals (Fig. 9J) express substantial levels of DARC and CD35 compared to RBCs.

**Fig. 9.**
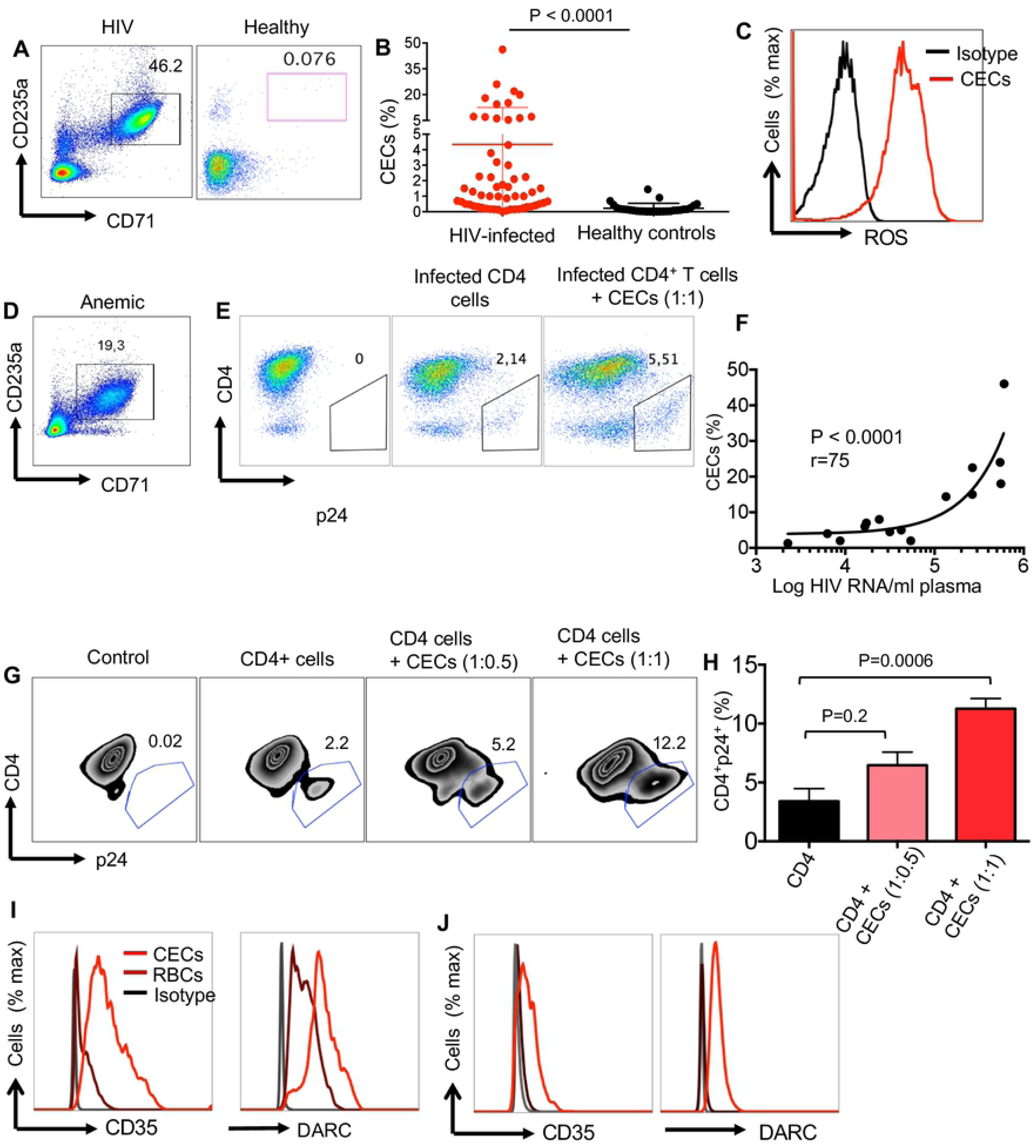
CECs from the peripheral blood of HIV-1 infected and anemic individuals enhance HIV infection in CD4^+^ T cells. (**A**) Representative flow cytometry plots showing frequency of CECs in an HIV-infected individual versus healthy control. (**B**) Cumulative data indicating %CECs in HIV-infected versus healthy individuals. (**C**) Representative histogram showing ROS expression in CECs of an HIV-patient (red line) and isotype control (black line). (**D**) Representative flow cytometry plots showing frequency of CECs in an anemic individual. (**E**) Representative flow cytometry plots showing HIV-infection in activated CD4^+^ T cells alone or in the presence of autologous CECs obtained from an HIV-infected individual. (**F**) Data showing correlation of CECs with the plasma viral load in HIV-infected but ART-naïve individulas. (**G**) Representative flow cytometry plots showing HIV-infection in activated CD4^+^ T cells alone or in the presence of autologous CECs obtained from an anemic individual. (**H**) Cumulative data showing %CD4^+^p24^+^ T cells in the absence or presence of CECs from anemic individuals in a dose dependent manner. (**I**) Representative histogram showing CD35 and DARC expression in CECs (red line), RBCs (brown line) and isotype control (black line) from an HIV-infected individual. (**J**) Representative histogram showing CD35 and DARC expression in CECs (red line), RBCs (brown line) and isotype control (black line) from an anemic individual.

## Discussion

The association of HIV-1 with cell-surface markers on erythrocytes has not been free of controversy. Erythrocytes from HIV-1 infected individuals were reported to contain cell-bound virus, and proposed to represent as a novel HIV-1 reservoir (32). However, this concept was later disputed and the reservoir hypothesis for erythrocytes was challenged (33). Subsequent studies confirmed that CR1 (CD35) and DARC were responsible for HIV-1 binding to the erythrocytes and trans-infection of the virus to target cells (16, 31). Despite these reports, the interaction of erythroid precursors with HIV-1 has never been investigated and has remained a mystery. We have recently reported abundance of erythroid precursors (CECs) in the cord blood, placental tissues and the peripheral blood of pregnant mothers(10, 15, 34). We further indicated that CECs possess a wide range of immunological properties (35). Here we reveal a novel role for CECs-mediated enhanced HIV-1 infection/replication in CD4^+^ T cells. We demonstrate that CECs from human cord blood or placental tissues in a dose-dependent manner exacerbate HIV-1, both R5 and X-tropic viral strains, replication/infection in autologous CD4^+^ T cells. Although this process was most marked for already activated CD4^+^ T cells, CECs significantly enhanced HIV-1 infection in non-activated CD4^+^ T cells, as well. Furthermore, we discovered that CECs do not require cell-cell interactions but via soluble factors enhance HIV-1 infection/replication in autologous CD4^+^ T cells. To understand the mechanism associated with these observations, we found that production of arginase-2 and TGF-β (25) by CECs did not enhance HIV-infection. Therefore, to determine the mechanism (s) associated with the enhanced HIV infection/replication, RNAseq analysis was performed. We found the transcriptome profile of infected CD4^+^ T co-cultured with CECs was clearly separated from the infected CD4^+^ T cells in the absence of CECs. Gene ontology of biological process for the transcriptome profile revealed upregulation of cellular response to oxygen-containing compounds and upregulation of NF-kB signaling in CD4^+^ cells when co-cultured with CECs. Our subsequent studies revealed that CECs has substantial levels of NOX2 mRNA, while other NOX paralogous (NOX 1, 3, 4, 5, DUOX1, and 2) were undetectable. Although endogenous ROS generation by RBCs has been documented (36), we observed that CECs have significantly higher ROS production capacity compared to their mature counterparts. More importantly, we observed that CECs release mitochondrial ROS, which its function can be abrogated by ROS-scavenger Apocynin but not by N-acetyl cysteine. Interestingly, we observed that CECs when obtained from the cord blood of mothers with IBD did not enhance HIV-1 infection in autologous CD4^+^ T cells because of their impaired ROS production ability (10).

Subsequently, we found elevated NF-kB gene expression in CD4^+^ T cells when co-cultured with CECs, suggesting enhanced NF-kB expression by ROS. A difficulty in establishing ROS signaling is that ROS often functions differently in a given pathway (e.g. upstream or downstream). This seems to be the case with regards to the role of ROS in NF-kB pathway. For example, ROS stimulates the NF-kB pathway in the cytoplasm but not in the nucleus (37). In this report we believe that ROS released by CECs directly activates NF-kB(38) and ROS production by CECs might be an essential step leading to NF-kB activation. In agreement, IKBKB gene was highly upregulated (> 8 folds) in HIV-1 infected CD4^+^ T cells when exposed to CECs compared to HIV-1 infected CD4^+^ T cells alone.

Activation of NF-κB is considered a crucial step in HIV-1 replication(39, 40). Therefore, we believe increased intracellular levels of NF-kB may permit high levels of HIV-1 gene expression and could thus provide a favorable environment for HIV replication in CD4^+^ T cells in the presence of ROS. In addition to NF-kB upregulation, a wide range of other genes were highly unregulated in CD4^+^ T cells when co-cultured with CECs. Among them tTg was the most upregulated gene, which may be a natural and non-specific mechanism of CD4^+^ T cells defense(41), aimed to reduce viral spreading.

Another important upregulated gene was AQP9, which its expression in PBMCs of HIV-infected individuals has already been reported(42). Although, its role in HIV infection is unknown, we suggest AQP9 may act as a ROS scavenger to prevent T cells apoptosis(27). Similarly, upregulation of MYOF gene in HIV-1-infected cells has been documented(43). We propose that MYOF participates in the repair of the plasma membrane following the budding of a multitude of HIV-1 virions, allowing the cell to live longer and to produce more viruses, which is the case in our study. ASAP1was another highly upregulated gene in HIV-1 infected CD4^+^ T cells when exposed to CECs. Although its role in HIV-1 infection is unknown, overexpression of ASAP1enhances cancer cell proliferation and invasion(44). There is a possibility that upregulation of ASAP-1 and TCL1 inhibit apoptosis(45) of HIV-1 infected CD4^+^ T cells, enabling them to produce more viral particles. In addition, we observed upregulation of Lck gene in CD4^+^ T cells exposed to CECs, which suggests Lck may facilitate HIV-1 assembly and release from the T cell plasma membrane as reduction of Lck has been reported to reduce HIV replication in CD4^+^ T cells(46). BCLAF1 was another highly upregulated gene in infected CD4^+^ T cells when co-cultured with CECs, which may act as a HIV-1 restricting factor as reported for human cytomegalovirus(47).

The role of LRRK2 gene in HIV-1 pathogenesis is unknown however it plays an important role against intracellular pathogens such as *Mycobacterium leprae*(48), *Salmonella Typhimurium*(49) and *Listeria monocytogenes*(50) via inflammasome activation(51). Thus, we suggest it may play a similar role against HIV-1 infection. Notably, IDO1 and KYNU genes were profoundly upregulated in CD4^+^ T cells when CECs were present. IDO-1 driven generation of kynurenine has immune regulatory function and IDO-1 appears to prevent viral protein production(52). In this context, we believe IDO-1 and KYNU may act to limit viral replication, which is indicative of the high viral production in CD4^+^ T cells co-cultured with CECs. Another important gene to mention was PogZ. Interaction of PogZ with HIV achieves efficient integration, which results in increased infectivity (53). Collectively, these data suggest that CECs via ROS production upregulate a wide range of genes and some of them explain enhanced viral infection/replication in CD4^+^ T cells.

Another aspect of our study was to determine whether CECs via binding to HIV-1 can trans-infect the virus to uninfected CD4^+^ T cells. Although the exact mechanism of binding of HIV-1 to RBCs is not fully understood, it is reported that HIV-1 binds to CR1 (31) and DARC on RBCs (16, 54). Binding of HIV to CR1/DARC demonstrates that HIV might exploit CR1/DARC to its advantage as a unique niche that enables viral survival and trans-infection to HIV-target cells. It was striking that both DARC and CR1 were highly expressed on CECs compared to their mature counterparts in terms of intensity and frequency. Thus, we speculated that these two receptors might come into play that together confer an accelerated rate of HIV trans-infection to CD4^+^ T cells. In fact, we observed that pre-incubation CECs with HIV-1 enables the virus to bind to these cells and subsequently CECs trans-infect HIV to CD4^+^ T cells. To establish that these effects were mediated specifically by DARC and/or CR1, we investigated whether CCL-5 a chemokine ligand for DARC or antibody against CR1 can compete for the adherence of HIV to DARC/CR1 on CECs. Despite the previous reports that RBC-bound HIV occurs predominantly via DARC(16, 54), we found that using rCCL-5 (even at 1000x higher concentration) failed to prevent adherence of HIV to CECs. Although, CECs have greater intensity for DARC on their surface and almost 100% of them express DARC compared to approximately 20% of RBCs, we suggest DARC might not be the dominant receptor for HIV. Our results thus appear to be compatible with the previous reports that have challenged binding of HIV to DARC as a crucial factor in HIV pathogenesis(55–57). Similarly, despite previous reports for the role of CR1 on RBCs for enhanced HIV infectivity to target cells(31), in our hands anti-CR1 antibody also failed to block trans-infection of HIV to CD4^+^ T cells by CECs. Our observations do not exclude DARC and CR1 as possible RBCs binding sites since a partial reduction in HIV trans-infection was occurred when combined targeted strategy was applied (rCCL-5 + anti-CR1). However, highly efficient transfer of HIV from CECs to CD4^+^ T cells in the presence of rCCL-5 and CR1 blockade suggests another potential HIV binding molecule on CECs. Our further studies revealed that HIV interacts with CD235a on CECs as reported for hepatitis A virus(58). This might be explained by significantly higher intensity of CD235a on CECs versus RBCs, which enables CECs to become efficient HIV carriers. Our observations suggest cluster formation of CD235a following interactions with HIV-1. It has been suggested that RBC glycoproteins (e.g. CD235a) may act as decoy receptors for pathogens (59). In this context, attachment of CECs to HIV may constitute an advantage for the virus to get a free ride to different anatomical locations. To determine whether the *in vitro* findings have any *in vivo* relevance, we studied the frequency of CECs in fresh PBMCs isolated from HIV-infected individuals.

Interestingly, we found significantly higher abundance of CECs in the peripheral blood of HIV infected individuals compared to healthy controls. In agreement with CECs isolated from the cord blood and placental tissues, CECs from HIV-infected individuals also express substantial levels of DARC and CR1 and subsequently enhance HIV-replication via ROS. Similar results were observed when frequency and functionality of CECs from anemic individuals was studied.

It appears that CECs regardless of their origin have similar immunological properties and adhere to HIV and enhance its infection/replication in CD4^+^ T cells *in vitro*. Although there is more HIV associated with white blood cells (e.g. CD4, monocytes and dendritic cells) the striking greater number of erythrocytes in the blood makes these cells a major source of viral RNA and subsequently transmissible virus (32, 60). A positive correlation between frequency of CECs with plasma viral load in ART-naïve patients suggest an important role for CECs in HIV pathogenesis. If erythrocyte-associated HIV through immune complexes serve as viral reservoir(32, 55, 60), greater expression of CR1, DARC and CD235a on CECs provide additional evidence that CECs might act more efficiently than their older siblings to carry the virus to different target cells within the body. Anemia is commonly reported among HIV-infected children underlying nutritional deficiencies and endemic parasitic infections, such as malaria and helminths, which lead to RBC destruction, or decreased production. Therefore, these conditions lead to the abundance of CECs in the periphery (35).

Subsequently, presence of CECs may enhance HIV replication in infected individuals or may enhance HIV infection/acquisition under certain circumstances. For instance, physiological abundance of CECs in the cord blood, placenta and peripheral blood of mothers during pregnancy may enhance mother to child HIV transmission (MTCT). Although, this hypothesis has not been tested in this study but can be suggested as potential mechanism of MTCT. In addition, extramedullary erythropoiesis under pathological conditions (e.g. pregnancy, chronic infections and cancer)(35) may also impact HIV infection/replication and should be taken in consideration.

Finally, presence of infectious viral particle on the surface or inside of CECs in the presence of Tenofovir demonstrates the potential role of CECs in HIV pathogenesis, which should be considered in treatment strategies.

We acknowledge several limitations to our study. Although we were able to find a correlation between % CECs in HIV-patients and plasma viral load in ART-naïve individuals, most of HIV-patients were on ART and had extremely low to undetectable viral load. Future studies targeting ART-naïve individuals will provide a better insight into the role of CECs in HIV-1 replication/transmission. Secondly, lack of access to ART-naïve and HIV-1 infected pregnant mothers prevented us to conclude if the frequency or functionality of CECs in the cord blood/placenta and mother’s blood impact vertical HIV-1 transmission. Moreover, unavailability of blocking antibodies against CD235a did not provide us the opportunity to block CD235a interactions with HIV *in vitro*. However, previous report for the interaction of CD235a with hepatitis A virus supports our observation (58). Future studies will also be needed to determine whether there is any association between the viral reservoirs in anemic with high percentages of CECs versus non-anemic HIV-1 infected individuals.

In summary, our results support a conceptual model that the interplay between extramedullary erythropoiesis and abundance of CECs in the periphery may influence HIV pathogenesis. More importantly, these findings provide a novel role for CECs-mediated HIV infection/replication by ROS. By extension, the role of various cell-surface receptors (e.g. DARC, CD35, CD235a) and HIV trans-infection by CECs in the presence of ART, could highlight the current limitations of ART and present future targets for therapy.

## Methods

### Human subjects

A total of 90 cord blood and placental tissues were obtained from full-term healthy deliveries for these studies and in some cases from subjects with inflammatory bowel disease (IBD). Blood samples from > 50 HIV+ individuals, 40 healthy controls and 6 anemic individuals were also obtained. Plasma viral load and clinical data were obtained from the HIV clinic (SH).

### Ethics statement

The appropriate Institutional Review Boards at the University of Alberta approved the studies. All study participants gave written informed consent to participate in this study (Protocol # Pro00056685, Pro00046080 and Pro00046064). All study participants were adults >20 years old.

### Cell isolation

Peripheral blood mononuclear cells (PBMCs) or Cord blood mononuclear cells (CBMC) were isolated by density gradient using Ficoll-Paque premium according to our previous reports (12, 61). Placental tissues were processed according to our reports elsewhere(10, 34). CECs were purified from CBMC, placental tissues or PBMCs according our previous reports (6, 15). CD4^+^ T cells were also isolated by negative selection using enrichment kit (STEM CELL Technologies) according to instruction’s manual. The purity of isolated cells was normally > 95% (SI Appendix, Fig. S2G and H).

### HIV-1 viral isolates

The CXCR4-utilizing isolate LAI and the primary CCR5-utilizing HIV-1 strain CSF-Jr were obtained from the AIDS Research and Reagent Program at the NIH. The HIV-1 with GFP marker was kindly provided by Dr. Christopher Power’s laboratory at the University of Alberta.

### *Ex vivo* HIV infection

Purified CD4^+^ T cells were infected with the LAI, CSF-Jr or eGFP viral isolates at multiplicity of infection (MOI) of 0.1 using magnetofection, as we have previously described elsewhere (23, 62). In some experiments, freshly isolated CD4^+^ T cells without prior activation were infected with the virus or isolated CD4^+^ T cells were first co-cultured with CECs for 24 hours and then infected with the virus.

A fixed number of infected CD4^+^ T cells (normally 10^6^/ ml) were co-cultured with different ratios of CECs in the presence/absence of L-arginine (1 to 4 mM), TGF-β inhibitor (10 μM), N-acetyl cysteine (NAC; 1mM), and apocynin (1 μM- 1mM) in 24 or 48 well plates for 4 days. In some experiments, CECs and CD4^+^ T cells were co-cultured in a trans-well system.

### Flow cytometry analysis

Cells were stained with anti-CD4 (Clone RPA-T4) and anti-CD3 (Clone SK7) conjugated antibodies (BD Bioscience). LIVE/DEAD® Fixable Aqua Dead Cell Stain Kit (ThermoFisher Scientific) was used to exclude dead cells. Cells were then fixed and permeabilized with cytofix/cytoperm (BD Bioscience) followed by intracellular staining with KC57-PE conjugated anti-p24 antibody (Beckman Coulter).

CR-1 (E11), DARC (2C3), CD71 (OX-26), CD235a (GA-R2), CCR5 (3A9) and CXCR4 (12G5) all were purchased from BD. ROS staining (Sigma) and MitoSOX red mitochondrial superoxide indicator (ThermoFisher Scientific) were performed per manufacturer’s protocols. Paraformaldehyde-fixed cells were acquired using a BD LSR Fortessa flow cytometer (BD Biosciences,) and analyzed with FlowJo (version 10) software.

### Trans-infect assays

Enriched CECs were incubated with HIV at MOI of 0.1 either with or without magnetofection as described previously. Then cells were washed extensively and co-cultured with either PHA-activated or inactivated CD4^+^ T cells. In some experiments, CECs were pre-incubated with human recombinant CCL5 (1-1000 ng/ml, R&D) and/or CD35 blocking antibody (5-20 μg/ml, clone J3D3, Beckman Coulter) for an hour prior to infection.

### Gene expression analysis

RNA extraction gene expression assays were performed according to our previous reports (10, 23, 62). To determine the expression levels of genes involved in ROS production, SYBR Green Master mix (Qiagen) was used targeting the following genes: NOX1 (QT00025585), NOX2 (CYBB, QT00029533), NOX3 (QT00044737), NOX4 (QT00057498), NOX5 (QT00021924), DUOX1 (QT00038346), and DUOX2 (QT00012236). TaqMan probes were used to detect expression levels of NF-kB (NFKBIA, Hs00355671-g1) and arginase-2 (Arg2, Hs00982833-ml). Beta-2 microglobulin (B2M, QT00088935) and beta actin (ACTB, Hs01060665_g1) for Qiagen and TaqMan chemistry, respectively, was used as reference genes. Each sample was run in duplicates and individual reactions contained 10 ng cDNA in a total reaction volume of 20 μL. The gene expression of the targeted genes was calculated by the 2^−ΔΔCt^ method, and specific mRNA levels were expressed as fold change over the uninfected condition. **RNA sequencing.** RNA libraries were constructed using the TruSeq RNA Library Prep kit v2 (Illumina) and then sequenced on a NextSeq 500 instrument (Illumina) and a 150bp paired-end protocol at an average depth ∼ 23.3 M reads per sample (The Applied Genomic Core), University of Alberta. Raw data was deposited in the SRA database of NCBI and is publicly available under accession number PRJNA529907.

FASTQ files were subjected to quality control trimming all bases with a Q score lower than 20 and trimmed reads with length shorted than 75 bp were discarded. Pseudo-aligments were conducted with Kallisto, with 100 permutations, to generate abundance estimates (counts). Raw counts from the sample reads were subjected to statistical analysis using the R package EdgeR (version 3.20.9). Raw counts were transformed into a differentially expressed genes (DEG) list and subjected to differential gene expression analysis. Significantly upregulated and downregulated genes were identified as having a False Discovery Rate of FDR <0.05 (SI Appendix data A).

For heatmapping analysis, significantly differentially expressed genes with a logFC < −2 and logFC > 2 with a count per-million (CPM) in at least two samples > 2 CPM. The genes were plotted with the package pheatmap using Euclidean clustering and Ward aggregation (SI Appendix data B).

Clusters identified by heatmapping were subjected to gene ontology analysis using the Biological Process Gene Ontology function on the gene ontology consortium website.

### Image stream analysis

An Amnis Image stream Mark II (EMD millipore) was used to collect at least 5000 images for each sample and each condition Analysis was performed by selecting fluorescence intensity for each targeted cell marker.

### Statistical Analysis

Statistical comparison between various groups was performed by the t-test using PRISM software. Also, differences were evaluated using One-Way ANOVA followed by Tukey’s test for multiple comparisons. Results are expressed as mean± SEM. P value <0.05 was considered as statistically significance.

## Acknowledgments

This work was supported by an Operating Grant: Innovative Biomedical and Clinical HIC/AIDS Research (370867) from the Canadian Institutes of Health Research (CIHR), a CIHR New Investigator Salary Award (360929) and a CIHR Foundation Scheme Grant (353953) (all to S.E.). In addition, this study was supported in part by an Innovation Grant from the Women and Children’s Health Research Institute (WCHRI to S.E.). The authors would like to thank the University of Alberta Faculty of Medicine and Dentistry’s The Applied Genomics Core for their assistance. Also, the University of Alberta Faculty of Medicine and Dentistry, Flow cytometry facility, which has received financial support from the faculty of Medicine and Dentistry and the Canadian Foundation for Innovation (CFI) awards to contributing investigators. We also thank HIV-infected individuals from the Northern Alberta HIV program, WCHRI for the cord blood and placenta tissues and healthy individuals for their contribution in this study.

## Author contributions

A.N. performed most of the experiments and wrote the methods. G.D. performed RNAseq analysis and conduced Image Stream. P.K. assisted in cell isolation and performed qPCR. S.S. obtained blood from HIV-patients, measured CECs in HIV and healthy controls. J.J assisted in conducting RNAseq and data analysis. S.H. contributed in HIV-patients recruitment. S.E. performed the initial studies and conceived the original idea, designed and supervised all of the research, assisted in data analysis and wrote the manuscript.

---

**Fig. S1.** (**A**) Cumulative data showing MFI of p24 in infected-CD4^+^ T cells in the absence or presence of CECs. (**B**) Representative Amnis Imagestream plots showing intensity of p24 in CD4^+^ T cells infected with HIV in the absence and presence of CECs. (**C**) Representative flow cytometry plot showing CXCR4 and CCR-5 expression on non-activated CD4^+^ T cells from the cord blood. (**D**) Hierarchical clustering on Euclidian distances showing different gene expression profile in HIV-infected CD4^+^ T cells in the presence or absence of CECs. (**E**) Principal component analysis (PCA) on the Euclidian distances between HIV-infected CD4^+^ T cells in the presence or absence of CECs. (**F**) Selected highly upregulated and downregulated genes in HIV-infected CD4^+^ T cells in the presence of CECs versus HIV-infected CD4^+^ T cells. (**G**) Representative plots showing purity of CD4^+^ T cells pre- and post-isolation. (**H**) Representative plots showing purity of CECs post-isolation.

**Fig. S2.** (**A**) Showing gene ontology of biological process of the transcriptome profile of co-cultured CD4^+^ T cells with CECs. (**B**) Cumulative data showing mRNA expression levels for arginase-2 (Arg-2) in the cord blood CECs from healthy and non-IBD versus Ulcerative colitis or Chron’s disease. (**C**) Cumulative data showing mRNA expression levels for arginase-2 (Arg-2) in the placenta CECs from healthy and non-IBD versus Ulcerative colitis or Chron’s disease. (**D**) Representative flow cytometry plots showing HIV-infection in CD4^+^ T cells in the presence/absence of CECs or following exposure of CECs to HIV in the presence of anti-CD35 using serum free culture media. (**E**) Cumulative data showing HIV-infection in CD4^+^ T cells in the presence/absence of CECs or following exposure of CECs to HIV in the presence of anti-CD35 using serum free culture media. (**F**) Representative flow cytometry plots showing HIV-infection in non-activated CD4^+^ T cells following cu-culture with HIV-exposed CECs.

**Data SI A**. The script showing gene expression analysis on raw counts (non-normalized) RNASeq libraries.

**Data SI B.** The script of heatmapping analysis of raw counts data from differentially expressed genes.

## References

1. Bianconi E, Piovesan A, Facchin F, Beraudi A, Casadei R, Frabetti F, et al. An estimation of the number of cells in the human body. Ann Hum Biol. 2013;40(6):463–71.

2. Kuhn V, Diederich L, Keller TCSt, Kramer CM, Luckstadt W, Panknin C, et al. Red Blood Cell Function and Dysfunction: Redox Regulation, Nitric Oxide Metabolism, Anemia. Antioxid Redox Signal. 2017;26(13):718–42.

3. Anderson HL, Brodsky IE, Mangalmurti NS. The Evolving Erythrocyte: Red Blood Cells as Modulators of Innate Immunity. Journal of immunology. 2018;201(5):1343–51.

4. Darbonne WC, Rice GC, Mohler MA, Apple T, Hebert CA, Valente AJ, et al. Red blood cells are a sink for interleukin 8, a leukocyte chemotaxin. The Journal of clinical investigation. 1991;88(4):1362–9.

5. Elahi S. New insight into an old concept: role of immature erythroid cells in immune pathogenesis of neonatal infection. Frontiers in immunology. 2014;5:376.

6. Dunsmore G, Bozorgmehr N, Delyea C, Koleva P, Namdar A, Elahi S. Erythroid Suppressor Cells Compromise Neonatal Immune Response against Bordetella pertussis. Journal of immunology. 2017.

7. Miller D, Romero R, Unkel R, Xu Y, Vadillo-Ortega F, Hassan SS, et al. CD71+ erythroid cells from neonates born to women with preterm labor regulate cytokine and cellular responses. J Leukoc Biol. 2018.

8. Namdar A, Koleva P, Shahbaz S, Strom S, Gerdts V, Elahi S. CD71+ erythroid suppressor cells impair adaptive immunity against Bordetella pertussis. Sci Rep. 2017;7(1):7728.

9. Mohanty JG, Nagababu E, Rifkind JM. Red blood cell oxidative stress impairs oxygen delivery and induces red blood cell aging. Front Physiol. 2014;5:84.

10. Dunsmore G, Koleva P, Ghobakhloo N, Sutton RT, Ambrosio L, Meng X, et al. Lower Abundance and Impaired Function of CD71+ Erythroid Cells in Inflammatory Bowel Disease Patients During Pregnancy. J Crohns Colitis. 2018.

11. Zhao L, He R, Long H, Guo B, Jia Q, Qin D, et al. Late-stage tumors induce anemia and immunosuppressive extramedullary erythroid progenitor cells. Nature medicine. 2018;24(10):1536–44.

12. Delyea C, Bozorgmehr N, Koleva P, Dunsmore G, Shahbaz S, Huang V, et al. CD71(+) Erythroid Suppressor Cells Promote Fetomaternal Tolerance through Arginase-2 and PDL-1. Journal of immunology. 2018.

13. Stachon A, Segbers E, Holland-Letz T, Kempf R, Hering S, Krieg M. Nucleated red blood cells in the blood of medical intensive care patients indicate increased mortality risk: a prospective cohort study. Crit Care. 2007;11(3):R62.

14. Inra CN, Zhou BO, Acar M, Murphy MM, Richardson J, Zhao Z, et al. A perisinusoidal niche for extramedullary haematopoiesis in the spleen. Nature. 2015;527(7579):466–71.

15. Elahi S, Ertelt JM, Kinder JM, Jiang TT, Zhang X, Xin L, et al. Immunosuppressive CD71+ erythroid cells compromise neonatal host defence against infection. Nature. 2013;504(7478):158–62.

16. He W, Neil S, Kulkarni H, Wright E, Agan BK, Marconi VC, et al. Duffy antigen receptor for chemokines mediates trans-infection of HIV-1 from red blood cells to target cells and affects HIV-AIDS susceptibility. Cell host & microbe. 2008;4(1):52–62.

17. Mphatswe W, Blanckenberg N, Tudor-Williams G, Prendergast A, Thobakgale C, Mkhwanazia N, et al. High frequency of rapid immunological progression in African infants infected in the era of perinatal HIV prophylaxis. Aids. 2007;21(10):1253–61.

18. Muenchhoff M, Prendergast AJ, Goulder PJ. Immunity to HIV in Early Life. Frontiers in immunology. 2014;5:391.

19. Biberfeld G, Biberfeld P, Buonaguro F, Charpak N, de The G, Rea MF, et al. Mother-to-child transmission of HIV-1 - Meeting of World Federation of Scientists in Erice, Italy, August 2001. Joint Working Group - Report of AIDS and Infectious Diseases PMP, and Mother and Child Health PMP - Plea for action with special emphasis on antiretroviral therapy: a scientific and community challenge. Acta Paediatr. 2001;90(11):1337–9.

20. Kallianpur AR, Wang Q, Jia P, Hulgan T, Zhao Z, Letendre SL, et al. Anemia and Red Cell Indices Predict HIV-Associated Neurocognitive Impairment in the HAART Era. The Journal of infectious diseases. 2015.

21. Redig AJ, Berliner N. Pathogenesis and clinical implications of HIV-related anemia in 2013. Hematology Am Soc Hematol Educ Program. 2013;2013:377–81.

22. Wumba RD, Zanga J, Aloni MN, Mbanzulu K, Kahindo A, Mandina MN, et al. Interactions between malaria and HIV infections in pregnant women: a first report of the magnitude, clinical and laboratory features, and predictive factors in Kinshasa, the Democratic Republic of Congo. Malaria J. 2015;14.

23. Elahi S, Niki T, Hirashima M, Horton H. Galectin-9 binding to Tim-3 renders activated human CD4+ T cells less susceptible to HIV-1 infection. Blood. 2012;119(18):4192–204.

24. Cloke TE, Garvey L, Choi BS, Abebe T, Hailu A, Hancock M, et al. Increased Level of Arginase Activity Correlates with Disease Severity in HIV-Seropositive Patients. Journal of Infectious Diseases. 2010;202(3):374–85.

25. Shahbaz S, Bozorgmehr N, Koleva P, Namdar A, Jovel J, Fava RA, et al. CD71+VISTA+ erythroid cells promote the development and function of regulatory T cells through TGF-beta. PLoS Biol. 2018;16(12):e2006649.

26. Bernatchez PN, Sharma A, Kodaman P, Sessa WC. Myoferlin is critical for endocytosis in endothelial cells. Am J Physiol-Cell Ph. 2009;297(3):C484–C92.

27. Miki A, Kanamori A, Negi A, Naka M, Nakamura M. Loss of aquaporin 9 expression adversely affects the survival of retinal ganglion cells. The American journal of pathology. 2013;182(5):1727–39.

28. Fruehauf JP, Meyskens FL, Jr. Reactive oxygen species: a breath of life or death? Clin Cancer Res. 2007;13(3):789–94.

29. Gough DR, Cotter TG. Hydrogen peroxide: a Jekyll and Hyde signalling molecule. Cell Death Dis. 2011;2:e213.

30. Lambeth JD. NOX enzymes and the biology of reactive oxygen. Nature reviews Immunology. 2004;4(3):181–9.

31. Horakova E, Gasser O, Sadallah S, Inal JM, Bourgeois G, Ziekau I, et al. Complement mediates the binding of HIV to erythrocytes. Journal of immunology. 2004;173(6):4236–41.

32. Hess C, Klimkait T, Schlapbach L, Del Zenero V, Sadallah S, Horakova E, et al. Association of a pool of HIV-1 with erythrocytes in vivo: a cohort study. Lancet. 2002;359(9325):2230–4.

33. Fierer DS, Vargas J, Jr., Patel N, Clover G. Absence of erythrocyte-associated HIV-1 in vivo. The Journal of infectious diseases. 2007;196(4):587–90.

34. Delyea C, Bozorgmehr N, Koleva P, Dunsmore G, Shahbaz S, Huang V, et al. CD71(+) Erythroid Suppressor Cells Promote Fetomaternal Tolerance through Arginase-2 and PDL-1. Journal of immunology. 2018;200(12):4044–58.

35. Elahi S. Neglected Cells: Immunomodulatory Roles of CD71(+) Erythroid Cells. Trends Immunol. 2019.

36. George A, Pushkaran S, Konstantinidis DG, Koochaki S, Malik P, Mohandas N, et al. Erythrocyte NADPH oxidase activity modulated by Rac GTPases, PKC, and plasma cytokines contributes to oxidative stress in sickle cell disease. Blood. 2013;121(11):2099–107.

37. Kabe Y, Ando K, Hirao S, Yoshida M, Handa H. Redox regulation of NF-kappa B activation: Distinct redox regulation between the cytoplasm and the nucleus. Antioxid Redox Sign. 2005;7(3-4):395–403.

38. Van Antwerp DJ, Martin SJ, Verma IM, Green DR. Inhibition of TNF-induced apoptosis by NF-kappa B. Trends in Cell Biology. 1998;8(3):107–11.

39. Beauparlant P, Kwon H, Clarke M, Lin R, Sonenberg N, Wainberg M, et al. Transdominant mutants of I kappa B alpha block Tat-tumor necrosis factor synergistic activation of human immunodeficiency virus type 1 gene expression and virus multiplication. Journal of virology. 1996;70(9):5777–85.

40. Mhashilkar AM, Biswas DK, LaVecchio J, Pardee AB, Marasco WA. Inhibition of human immunodeficiency virus type 1 replication in vitro by a novel combination of anti-Tat single-chain intrabodies and NF-kappa B antagonists. Journal of virology. 1997;71(9):6486–94.

41. Amendola A, Rodolfo C, Di Caro A, Ciccosanti F, Falasca L, Piacentini M. “Tissue” transglutaminase expression in HIV-infected cells: an enzyme with an antiviral effect? Annals of the New York Academy of Sciences. 2001;946:108–20.

42. Zhao F, Ma JM, Huang LH, Deng Y, Li LQ, Zhou Y, et al. Comparative transcriptome analysis of PBMC from HIV patients pre- and post-antiretroviral therapy. Meta Gene. 2017;12:50–61.

43. Imbeault M, Giguere K, Ouellet M, Tremblay MJ. Exon level transcriptomic profiling of HIV-1-infected CD4(+) T cells reveals virus-induced genes and host environment favorable for viral replication. PLoS Pathog. 2012;8(8):e1002861.

44. Zhang T, Zhao GN, Yang CH, Dong PX, Watari H, Zeng L, et al. Lentiviral vector mediated-ASAP1 expression promotes epithelial to mesenchymal transition in ovarian cancer cells. Oncol Lett. 2018;15(4):4432–8.

45. Weng J, Rawal S, Chu F, Park HJ, Sharma R, Delgado DA, et al. TCL1: a shared tumor-associated antigen for immunotherapy against B-cell lymphomas. Blood. 2012;120(8):1613–23.

46. Strasner AB, Natarajan M, Doman T, Key D, August A, Henderson AJ. The Src kinase Lck facilitates assembly of HIV-1 at the plasma membrane. Journal of immunology. 2008;181(5):3706–13.

47. Lee SH, Kalejta RF, Kerry J, Semmes OJ, O’Connor CM, Khan Z, et al. BclAF1 restriction factor is neutralized by proteasomal degradation and microRNA repression during human cytomegalovirus infection. Proceedings of the National Academy of Sciences of the United States of America. 2012;109(24):9575–80.

48. Zhang FR, Huang W, Chen SM, Sun LD, Liu H, Li Y, et al. Genomewide association study of leprosy. The New England journal of medicine. 2009;361(27):2609–18.

49. Gardet A, Benita Y, Li C, Sands BE, Ballester I, Stevens C, et al. LRRK2 is involved in the IFN-gamma response and host response to pathogens. Journal of immunology. 2010;185(9):5577–85.

50. Zhang Q, Pan Y, Yan R, Zeng B, Wang H, Zhang X, et al. Commensal bacteria direct selective cargo sorting to promote symbiosis. Nature immunology. 2015;16(9):918–26.

51. Liu W, Liu X, Li Y, Zhao J, Liu Z, Hu Z, et al. LRRK2 promotes the activation of NLRC4 inflammasome during Salmonella Typhimurium infection. The Journal of experimental medicine. 2017;214(10):3051–66.

52. Kane M, Zang TM, Rihn SJ, Zhang F, Kueck T, Alim M, et al. Identification of Interferon-Stimulated Genes with Antiretroviral Activity. Cell host & microbe. 2016;20(3):392–405.

53. Tesina P, Cermakova K, Horejsi M, Prochazkova K, Fabry M, Sharma S, et al. Multiple cellular proteins interact with LEDGF/p75 through a conserved unstructured consensus motif. Nature communications. 2015;6.

54. Lachgar A, Jaureguiberry G, Le Buenac H, Bizzini B, Zagury JF, Rappaport J, et al. Binding of HIV-1 to RBCs involves the Duffy antigen receptors for chemokines (DARC). Biomed Pharmacother. 1998;52(10):436–9.

55. Beck Z, Brown BK, Wieczorek L, Peachman KK, Matyas GR, Polonis VR, et al. Human Erythrocytes Selectively Bind and Enrich Infectious HIV-1 Virions. PloS one. 2009;4(12).

56. Horne KC, Li X, Jacobson LP, Palella F, Jamieson BD, Margolick JB, et al. Duffy Antigen Polymorphisms Do Not Alter Progression of HIV in African Americans in the MACS Cohort. Cell host & microbe. 2009;5(5):415–7.

57. Walley NM, Julg B, Dickson SP, Fellay J, Ge D, Walker BD, et al. The Duffy Antigen Receptor for Chemokines Null Promoter Variant Does Not Influence HIV-1 Acquisition or Disease Progression. Cell host & microbe. 2009;5(5):408–10.

58. Sanchez G, Aragones L, Costafreda MI, Ribes E, Bosch A, Pinto RM. Capsid region involved in hepatitis A virus binding to glycophorin A of the erythrocyte membrane. Journal of virology. 2004;78(18):9807–13.

59. Gagneux P, Varki A. Evolutionary considerations in relating oligosaccharide diversity to biological function. Glycobiology. 1999;9(8):747–55.

60. Levy JA. HIV-1: hitching a ride on erythrocytes. Lancet. 2002;359(9325):2212–3.

61. Elahi S, Dinges WL, Lejarcegui N, Laing KJ, Collier AC, Koelle DM, et al. Protective HIV-specific CD8+ T cells evade Treg cell suppression. Nature medicine. 2011;17(8):989–95.

62. Elahi S, Weiss RH, Merani S. Atorvastatin restricts HIV replication in CD4+ T cells by upregulation of p21. Aids. 2016;30(2):171–83.

